# The genome of the toxic invasive species *Heracleum sosnowskyi* carries an increased number of genes despite absence of recent whole-genome duplications

**DOI:** 10.1101/2023.02.14.528432

**Authors:** MI Schelkunov, VYu Shtratnikova, AV Klepikova, MS Makarenko, DO Omelchenko, LA Novikova, EN Obukhova, VP Bogdanov, AA Penin, MD Logacheva

## Abstract

*Heracleum sosnowskyi*, belonging to a group of giant hogweeds, is a plant with large effects on ecosystems and human health. It is an invasive species that contributes to the deterioration of grassland ecosystems. The ability of *H. sosnowskyi* to produce linear furanocoumarins (FCs), photosensitizing compounds, makes it very dangerous. At the same time, linear FCs are compounds with high pharmaceutical value that are used in skin disease therapies. Despite this high importance, it has not been the focus of genetic and genomic studies. Here, we report a chromosome-scale assembly of the Sosnowsky’s hogweed genome. Genomic analysis revealed an unusually high number of genes (55 206) in the hogweed genome, in contrast to the 25-35 thousand found in most plants. However, we did not find any traces of recent whole genome duplications not shared with its confamiliar, *Daucus carota* (carrot), which has approximately thirty thousand genes. The analysis of the genomic proximity of duplicated genes indicates tandem duplications as a main reason for this increase. We performed a genome-wide search of the genes of the FC biosynthesis pathway and their expression in aboveground plant parts. Using a combination of expression data and phylogenetic analysis, we found candidate genes for psoralen synthase and experimentally showed the activity of one of them using a heterologous yeast expression system. These findings expand our knowledge on the evolution of gene space in plants and lay a foundation for further analysis of hogweed as an invasive plant and as a source of FCs.

## Introduction

The immense influence of plants on the environment and human activity is not limited to beneficial effects. Many plants produce secondary metabolite compounds, which are toxic to people and animals. Some plants can greatly increase their area of distribution and transfer to new areas, thus causing threats to native ecosystems and human activity (invasive species). Notorious examples of invasive plant species that cause serious ecological and socioeconomic consequences are water hyacinth (Villamagna and Murphy, 2010), kudzu vine (Forseth and Innis, 2004), and Japanese knotweed (Lavoie, 2017). The object of our study, *Heracleum sosnowskyi* Manden. (Sosnowsky’s hogweed), possesses both of these characteristics. *H. sosnowskyi*, as well as its close relative *H. mantegazzianum* Sommier & Levier (both native to the Caucasus mountains), is a highly aggressive invasive plant that greatly increased its distribution area in Eastern and Western European countries. Both species have very large biomass, growth speed, and seed productivity (Figure 1A); this allows them to successfully increase their presence, leading to a decrease in species richness in local plants, soil insects and microorganisms, which they affect by releasing specific chemicals (Renčo and Baležentiené, 2015; Čerevková *et al.*, 2020), reviewed in (Grzędzicka, 2022). According to a recent forecast, ongoing climate change will allow Sosnowsky’s hogweed to greatly increase its distribution area in the next 20-40 years (Koldasbayeva *et al.*,2022). In addition to its adverse role in ecosystems, Sosnowsky’s hogweed is notorious for causing severe skin irritation and burns due to the presence of furanocoumarins (FCs), which are photosensitizing compounds (Jakubowicz *et al.*, 2012). Linear FCs, such as psoralen and its derivatives, are strong photosensitizers, so contact with hogweed and subsequent exposure to sunlight leads to serious skin damage (Rzymski *et al.*, 2015). On the other hand, psoralen and metoxy-psoralens are an integral part of therapy (psoralen UVA, PUVA therapy) for psoriasis and some other skin diseases (Armstrong and Read, 2020). Thus, hogweed can be used as a source of pharmaceutically relevant FCs. The insights into both of these aspects of *H. sosnowskyi* - its success as an invasive plant and its ability to synthesize linear FCs - require a solid genetic basis. In particular, the resolution of a so-called paradox of invasive species (Estoup *et al.*, 2016) in the case of Sosnowsky’s hogweed could have involved hybridization between *H. sosnowskyi* and related species (in particular, with the very close congener *H. mantegazzianum)* (Jahodová *et al.*, 2007).

**Figure 1.**
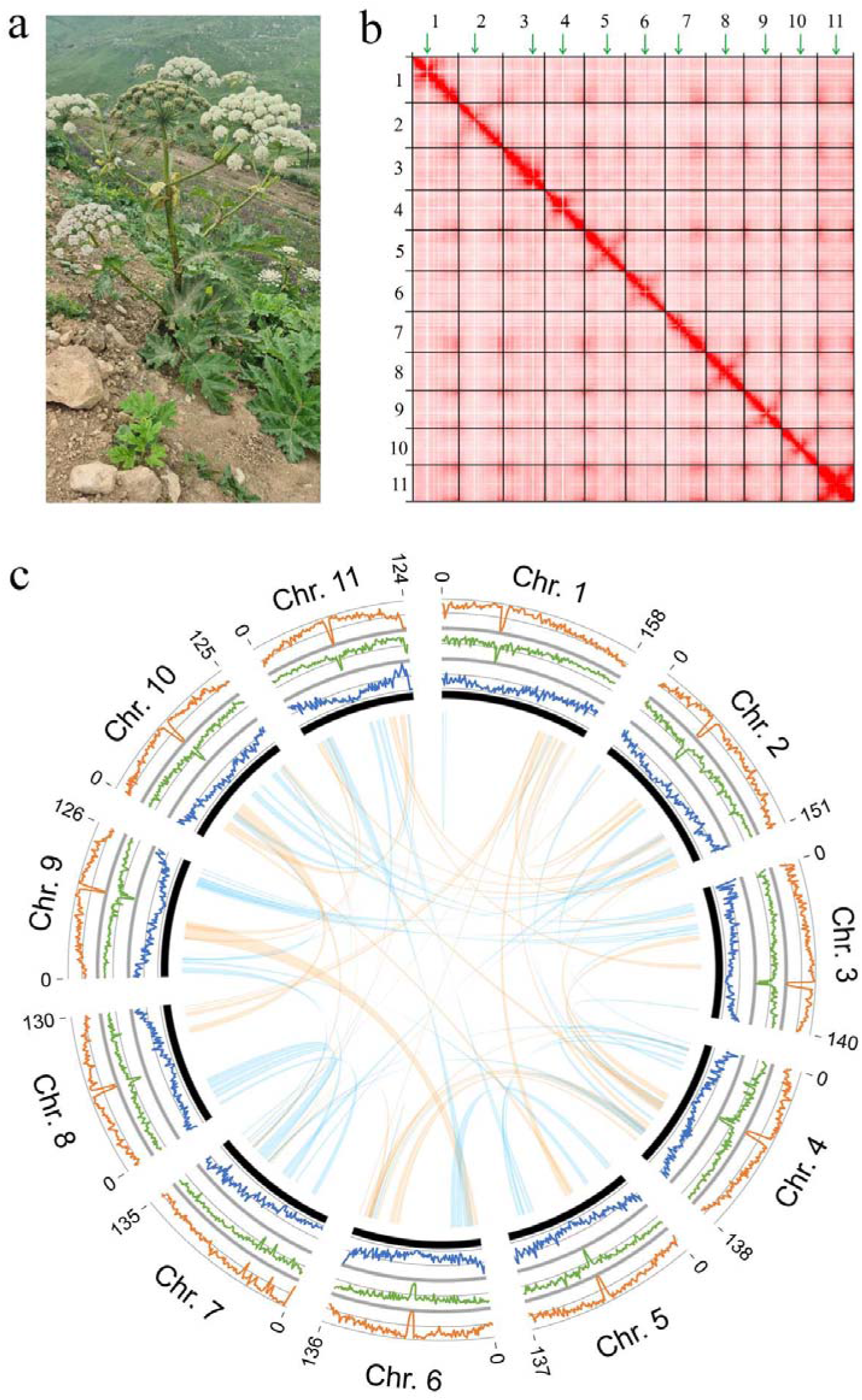
a: *Heracleum sosnowskyi*; general view of a plant. b: HiC-based map of chromatin contacts in the *H. sosnowskyi* genome assembly (only the 11 largest scaffolds corresponding to chromosomes are shown). Green arrows indicate centromeric repeats. c: Genome diagram for 11 *H. sosnowskyi* chromosomes. Cyan ribbons denote collinear blocks in direct orientation, orange – in reverse orientation. Orange lines outside of the circle denote the fraction of transposons in 1-Mb windows, green lines – gene expression level, blue lines – gene density (number of genes per megabase).

In addition to the immediate importance of this species, it belongs to the family Apiaceae, a large and underexplored group of plants. This family includes many economically important plants (carrot, parsley, celery, anise, and fennel), but high-quality genome-wide data are lacking; chromosome-scale assemblies are published only for few agricultural plants - carrot (Iorizzo *et al.*, 2016), coriander (Song *et al.*, 2020) and celery (Song *et al.*, 2021) and a medicinal plant *Angelica sinensis* (Han *et al.*, 2022). The genomes of Apiaceae are quite large (mean 2.18 Gb, according to (Leitch *et al.*, 2019)) and complex having undergone one or several whole genome duplications in a common ancestor of the family. The rapid advances in DNA sequencing technologies have equipped researchers with tools capable of tackling very complex eukaryotic genomes. In particular, HiFi (CCS) reads from the Pacific Bioscience platform coupled with chromatin conformation capture data have become the gold standard for plant genomes. Thus, we assembled the Sosnowsky’s hogweed genome and performed gene analysis focusing on the furanocoumarin biosynthesis pathway.

## Results and discussion

### Genome assembly and annotation

As a basis for the assembly, we used data from the Pacific Bioscience platform (1.4 million circular consensus reads with a median length of 13 091 bp). We tested several genome assemblers optimized for long reads (Hifiasm, Flye, Canu, and Miniasm) and inspected metrics such as total assembly size, N50 and the fraction of complete BUSCO genes (Supplementary Table S2). We found that Hifiasm performed better than other assemblers; however, it produced many duplicated genes due to incomplete collapsing of haplotypes. This is a well-known problem in eukaryotic genome assembly; recently, it was shown that it may affect hundreds and thousands of genes, leading to false biological conclusions (Ko *et al.*, 2022). To overcome this problem, we used the tool Mabs. Mabs is an add-on to long-read assemblers that allows automatic optimization of assembly parameters (Schelkunov, 2022). The application of Mabs improved the assembly both in terms of N50 and a fraction of duplicate genes (Table 1, Supplementary Table A2). The inspection of coverage distribution for BUSCO genes shows that duplicate genes retained by Mabs have the same coverage as single-copy ones, while raw Hifiasm assembly results in a two-peaked distribution where the peak with lower coverage corresponds to uncollapsed haplotypes (Supplementary Figure S1). We then used Mabs assembly for scaffolding with HiC data; the resulting scaffolds had an N50 of 137 Mb (Table 1). After manual correction based on the examination of HiC contact maps, we obtained 11 scaffolds that contained 92.4% of the total assembly length. Given that the haploid chromosome number in hogweed is 11 (Bell and Constance, 1966), we assumed that these scaffolds correspond to the 11 chromosomes of *H. sosnowskyi.* All these scaffolds carry a specific type of tandem repeat that is localized in one narrow region of the scaffold and not found outside of this region. This region presumably corresponds to a centromere and a repeat - to centromeric repeat (Figure 1B). We also looked for a telomeric repeat, which has a conserved DNA sequence, and found it at the ends of some (but not all) large scaffolds. This indicates high contiguity of the assembly, but there is also potential for improvement in telomeric and subtelomeric regions. To assess the completeness of the assembly in terms of gene content, we calculated the BUSCO score. Depending on the set of BUSCO genes (Viridiplantae or eudicots), the fraction of complete BUSCO genes ranged from 98.4 to 99.3% (Table 1). These values were similar to or higher than those for genomes of other species of Apiaceae (Song *et al.*, 2021), indicating a high level of completeness. The plastid genome of *H. sosnowskyi* is 149 368 bp in length, and it has the typical structure and gene content found in other representatives of Apiaceae. Similar to other species of the tribe Tordylieae, the *H. sosnowskyi* plastome has a reduced inverted repeat (IR) at the border with a large single copy region (LSC). In contrast to most plants that have IR, including *ycf2*, *trnI-CAU*, *rpl2* and *rpl23*, in *H. sosnowskyi*, these genes make up a part of the LSC (Supplementary Figure S2). Whole plastome phylogenetic analysis shows that *H. sosnowskyi* is grouped with other species of the genus and with related monotypic genera *Symphyoloma* and *Mandenovia* (Supplementary Figure S3). The same was inferred from the analysis of nuclear markers (Logacheva *et al.*, 2010); the results are also congruent with those of a plastid phylogenomic study that included species of *Heracleum* other than *H. sosnowskyi* (Samigullin *et al.*, 2022).

**Table 1.** Information on nuclear contigs and scaffolds. Plastid sequences, mitochondrial sequences and sequences of contamination are not counted. The difference in total length of scaffolds and contigs is non-informative since all gaps in these scaffolds consist of 200 “N” letters. C means complete BUSCO genes, S – single copy, D – duplicated, F – fragmented and M – missing.

|  | *Nuclear contigs* | *Nuclear scaffolds* |
| --- | --- | --- |
| Total length (bp) | 1 629 156 283 | 1 629 177 483 |
| N50 (bp) | 22 773 521 | 136 614 956 |
| Number of sequences | 489 | 383 |
| Longest sequence | 64 363 759 | 158 968 167 |
| Shortest sequence | 9 633 | 9 633 |
| BUSCO results by the "Eudicots" dataset | C: 98.4% [S: 89.8%, D: 8.6%], F: 0.2%, M: 1.4%, n: 2326 | C: 98.4% [S: 89.8%, D: 8.6%], F: 0.2%, M: 1.4%, n: 2326 |
| BUSCO results by the "Viridiplantae" dataset | C: 99.3% [S: 92.7%, D: 6.6%], F: 0.2%, M: 0.5%, n: 425 | C: 99.3% [S: 92.9%, D: 6.4%], F: 0.2%, M: 0.5%, n: 425 |

The annotation of the repeats show that they constitute 78.7% of the hogweed genome. This is similar to other Apiaceae species with large genome sizes (Song *et al.*, 2021; Song *et al.*, 2020). Retroelements constitute a prevalent part of the *H. sosnowskyi* repeatome; among these, Ty1/Copia and Gypsy/DIRS1 are the most frequent (34.9 and 14.5% of all retroelements) (Supplementary Figure S4). The distribution of repeat elements across the chromosomes is uneven, with some regions of the scaffold having a higher density than the others. As mentioned above, there is a repeat that presumably corresponds to a centromeric repeat (Figure 1B); the regions surrounding the centromere are rich in repeats (Supplementary Figure S5), mostly due to the low-complexity tandem, while the mobile elements demonstrate the reverse pattern: they are depleted in the centromere (Figure 1C). The genes are confined to regions distant from the centromere. The cluster of rRNA genes was found on chromosome 7. The distribution of repeats and genes in hogweed is similar to that in the other Apiaceae genomes (Supplementary Figures S6, S7, S8), including carrot, regardless of whether different elements of the genome were investigated cytogenetically (for review see (Iovene and Grzebelus, 2019)). This supports the correspondence of the large scaffolds to chromosomes.

Our approach to the annotation of protein-coding genes combined the use of transcriptome data and homology-based prediction. Initially, 71 331 genes were predicted; 55 106 were retained for final annotation after filtering (Supplementary Figure S9). However, the number of genes is strikingly greater than that usually found in most plants that are not recent polyploids, particularly in Apiaceae (Iorizzo *et al.*, 2016; Song *et al.*, 2021; Song *et al.*,2020). This might stem from several reasons: 1) technical problems with the assembly and/or annotation of the hogweed genome, which can lead to overestimation of the gene number (e.g., uncollapsed haplotypes or the splitting of a single gene into several during the annotation); 2) taxon-specific expansions of gene families in hogweed; and 3) underestimation of the gene number in other Apiaceae species. To test for these hypotheses, we performed several additional analyses involving our own data and previously reported genomes. First, we estimated the distribution of average coverage of hogweed genes belonging to different groups relative to their copy number. There was no decrease in the coverage for two- and multicopy genes (Supplementary Figure S10). This demonstrated that the increase in gene number is not due to the presence of uncollapsed haplotypes. The average CDS length was similar to that in other plants (e.g., *Arabidopsis* - 1218.3 bp, tomato - 1041.8 bp, and maize - 1172.5 bp), the number of exons, and the fraction of single-exon genes (Table 2) indicated that the annotation artefacts (splitting of genes) were also unlikely to be the reason. Thus, the high number of genes is a biological phenomenon, not a technical artefact; to determine whether it is unique for hogweed, we performed reannotation of the coriander, celery, and carrot genomes using the same pipeline that we used for hogweed (see methods). The results showed that the number of genes in Apiaceae might have been greatly underestimated, except for carrot (Table 2). The average exon and intron lengths were very similar in all four species; carrot had slightly higher gene and CDS lengths (Table 2, Figure 2A), but this is caused by the underprediction of exons in hogweed, celery and coriander (these species have much larger genomes than carrot, which can make the prediction of short exons more challenging) than by the splitting of genes because the fraction of single-exon genes was similar in these species. To assess the quality of these new annotations compared to the initial annotations, we performed BUSCO analysis of the set of annotated genes. The new annotations had a much higher fraction of complete genes and a lower fraction of missing and fragmented genes than the initial annotations (Figure 2B; note that initial annotations were available only for carrot and coriander). Thus, we use them in all downstream comparative analyses.

**Table 2.**
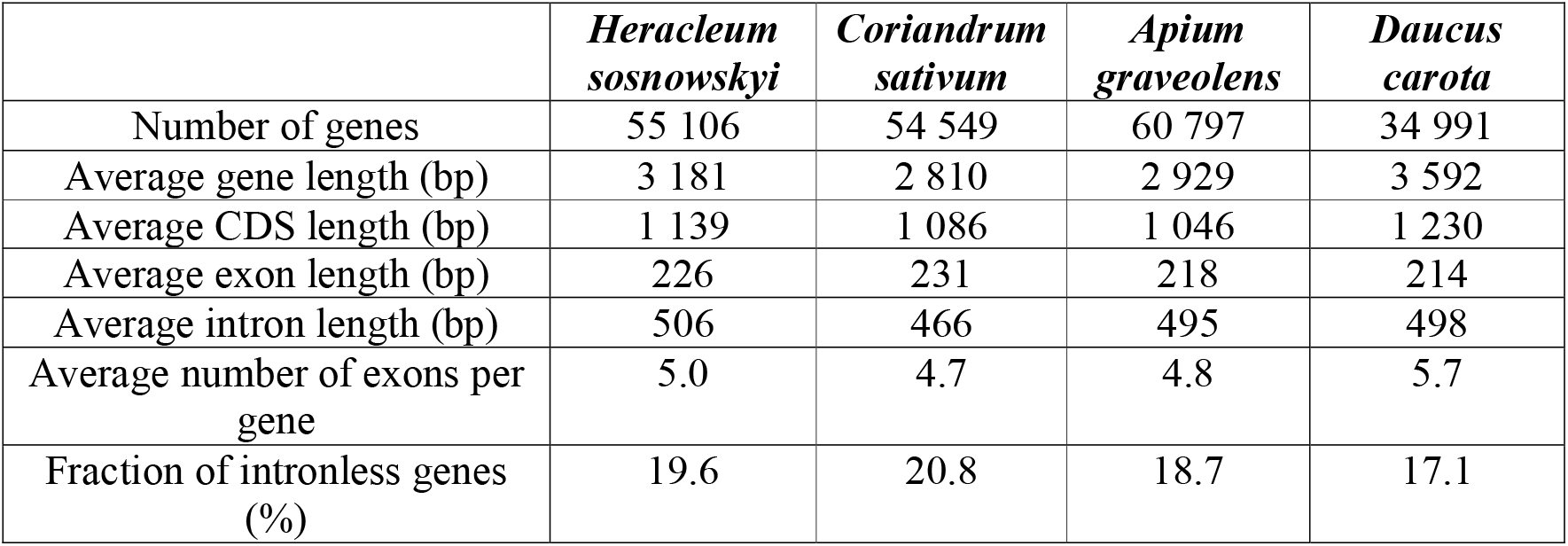
Statistics on protein-coding genes of *H. sosnowskyi* and other species of Apiaceae.

|  | *Heracleum sosnowskyi* | *Coriandrum sativum* | *Apium graveolens* | *Daucus carota* |
| --- | --- | --- | --- | --- |
| Number of genes | 55 106 | 54 549 | 60 797 | 34 991 |
| Average gene length (bp) | 3 181 | 2 810 | 2 929 | 3 592 |
| Average CDS length (bp) | 1 139 | 1 086 | 1 046 | 1 230 |
| Average exon length (bp) | 226 | 231 | 218 | 214 |
| Average intron length (bp) | 506 | 466 | 495 | 498 |
| Average number of exons per gene | 5.0 | 4.7 | 4.8 | 5.7 |
| Fraction of intronless genes (%) | 19.6 | 20.8 | 18.7 | 17.1 |

**Figure 2.**
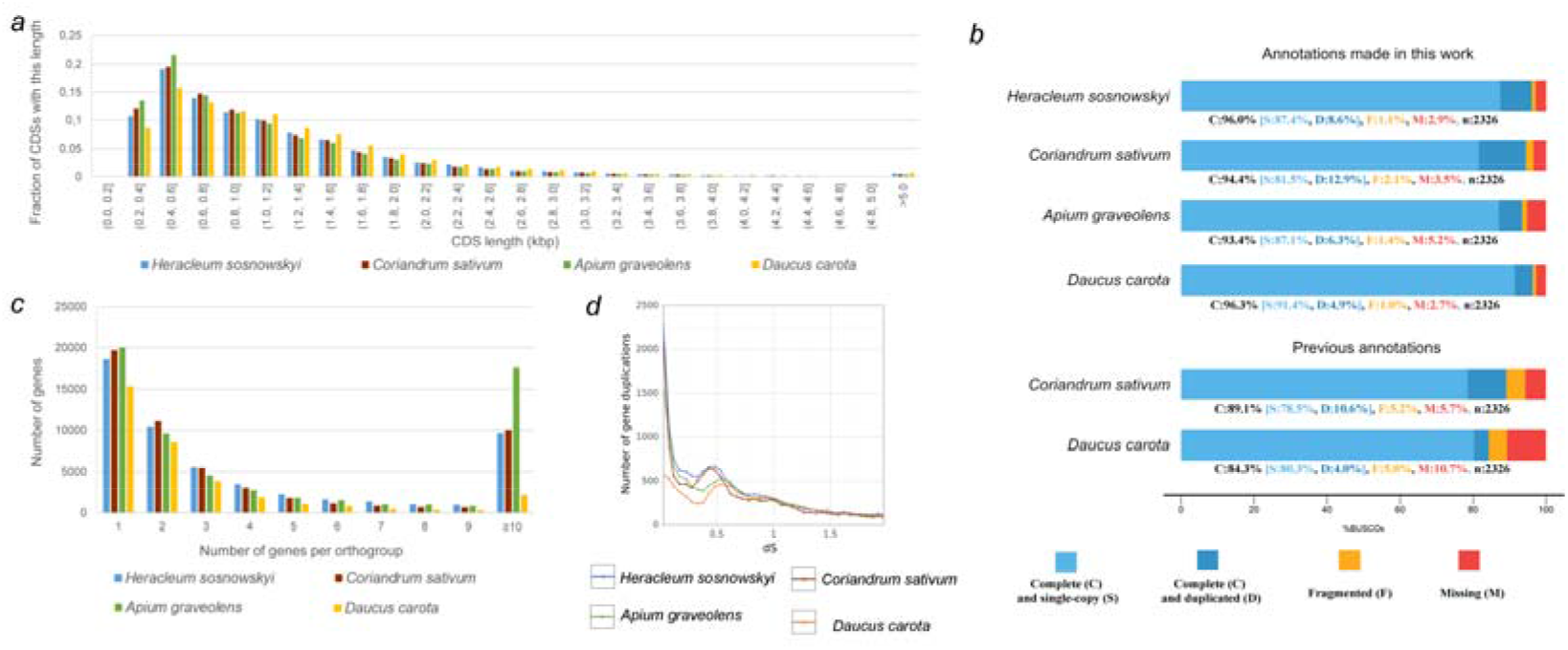
a: Distribution of CDS length in *H. sosnowskyi* and other species of Apiaceae b: Diagram showing BUSCO metrics for the genome annotations of *H. sosnowskyi* and other Apiaceae (upper panel – annotations produced in this study, lower panel – original annotations). c: Number of genes per orthogroup in *H. sosnowskyi* and other species of Apiaceae d: Distribution of dS (synonymous substitutions per synonymous site) between pairs of paralogues in *H. sosnowskyi* and other species of Apiaceae

### Expansion of gene families in the apioid superclade

As mentioned above, we found a striking increase in the number of genes in celery, coriander and hogweed compared to carrot. These three species are more closely related to one another, being the members of a group within Apiaceae that is called the **apioid superclade**, while carrot belongs to the outgroup tribe Daucinae (Downie *et al.*, 2001). The most obvious source of genes is whole genome duplication. According to earlier comparative genomic studies, an ancient WGD apparently occurred in a common ancestor of Apiaceae because it is shared between carrot, celery and coriander. This WGD is dated to 34-38 m.y.a.(Song *et al.*, 2021). Given the drastically different gene numbers in carrot and in the members of the apioid superclade, we propose three hypotheses: 1) a common ancestor of the apioid superclade underwent one or more additional WGD events that were not shared by carrot; 2) the common ancestor of the apioid superclade had a lower rate of genome fractionation (in particular, the loss of genes) after WGD compared to carrot; and 3) a higher number of genes is explained by other mechanisms not associated with WGD, for example, segmental duplications. We performed several analyses to test these hypotheses. First, we compared the size of orthogroups in carrot and in the members of the apioid superclade. There was no increase in the number of orthogroups that contained two genes, as expected for the WGD. Instead, there was an increase in the number of large orthogroups (10 and more genes) (Figure 2C). To gain further insight into the increase in gene number, we analysed the distribution of synonymous substitutions per synonymous site (dS), which is a reliable indicator of WGD (see, e.g., (Blanc and Wolfe, 2004), (Maere *et al.*, 2005)). The results showed no additional dS peaks in either hogweed or other members of the apioid superclade (Figure 2D). This is congruent with the results of (Song *et al.*, 2021), who reported no lineage-specific WGD in coriander and celery. The results of synteny analysis between hogweed and carrot do not support the hypothesis of additional WGD as well. In the case of WGD, one would expect the 2-to-1 correspondence of large genomic segments, which is not true, and 1-to-1 correspondence is prevalent (Figure 3A); the same is true for the comparison between carrot and other members of the apioid superclade (Supplementary Figure S11). To select between the other two hypotheses, we analysed the distribution of distances between paralogues that are present in the apioid superclade but not in carrot. Specifically, we considered the orthogroups that contained one carrot gene and two, three, four and five or more hogweed (or other apioid superclade species) genes. If the higher number of hogweed genes is explained by a lower rate of gene loss after WGD, those multiple genes are likely to be located on different chromosomes. If they are derived from smaller scale duplications (tandem or larger segmental), they are likely to be on one chromosome. Thus, we assessed genomic proximity between pairs of duplicated genes and compared it with that of randomly selected pairs of genes. For a random pair of genes, the probability of being on the same chromosome was ~0.1. The distance between random pairs of genes located on the same chromosome ranged from 49 to 111 Mb, depending on the species (Supplementary Table S3). For duplicated genes, both the fraction of genes located in one chromosome and the average distance between pairs of duplicates significantly differed from the randomly selected genes. The fraction of genes located in one chromosome differed from 0.23 to 0.53 depending on the size of the orthogroup. The average distance between pairs of duplicated genes that are on the same chromosome ranged from 3 to 17 Mb (Table 3). The distribution of distances showed that it is highly biased towards lower values (Figure 3B, C) for the orthogroups of all sizes (2, 3, 4 and 5 or more hogweed genes). We observed the same patterns in two other species of the apioid superclade (see Supplementary Tables S4 and S5 and Supplementary Figure S12). This high proximity of the duplicates indicated their origin by small-scale events, such as intrachromosomal segmental and tandem duplications.

**Figure 3.**
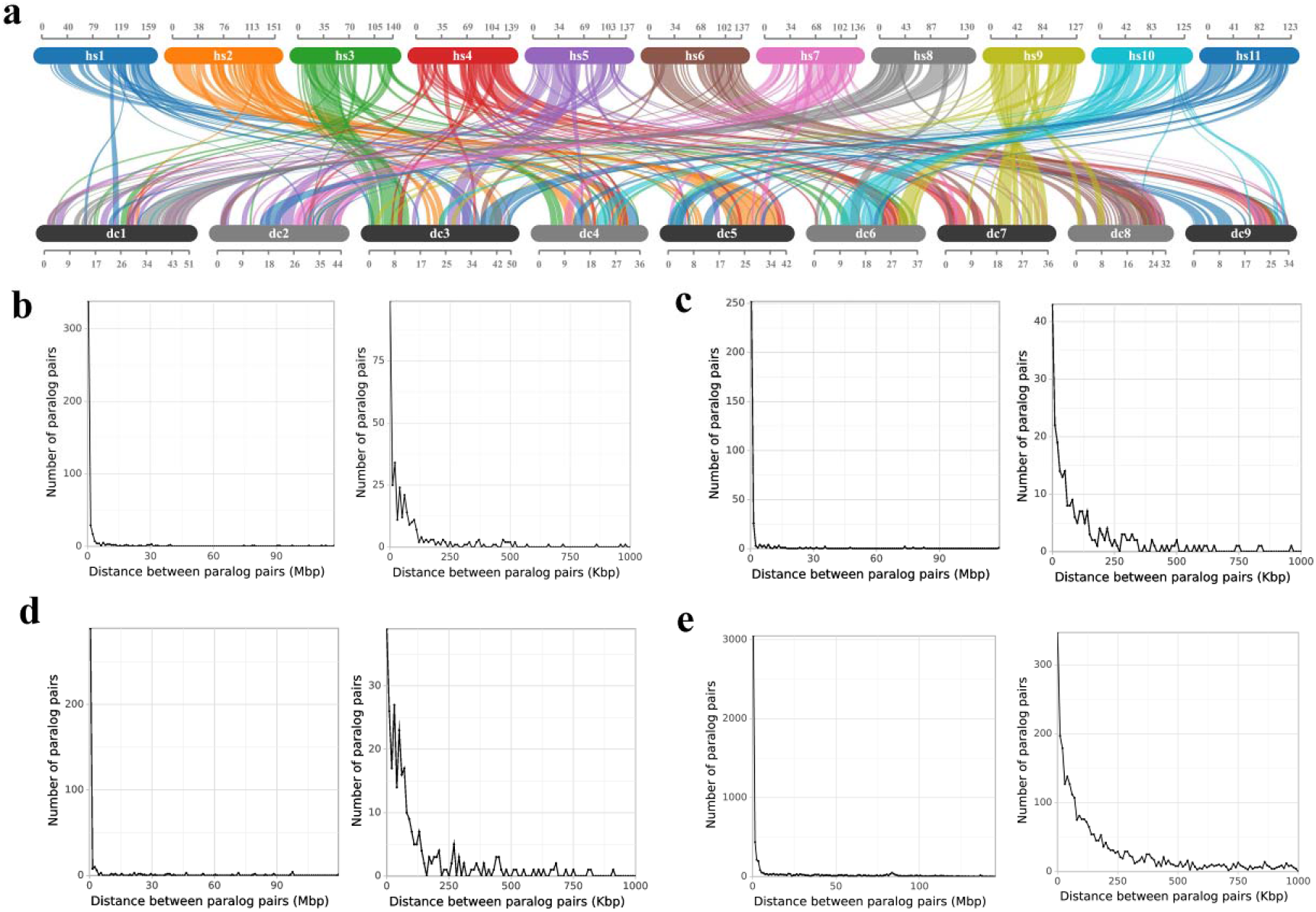
a: Collinear blocks between the genomes of *Heracleum sosnowskyi* (chromosomes denoted as hs1 - hs11) and *Daucus carota* (chromosomes denoted as dc1 – dc9). Only blocks that contain at least 20 pairs of homologous genes are shown. Coordinates are indicated in Mb. b: Distance between pairs of paralogues in orthogroups that have one *Daucus carota* and two *Heracleum sosnowskyi* genes, left – whole range of variation of distances, 1-Mb bins, right – distances up to 1 Mb, 10-Kb bins. c: Distance between pairs of paralogues in orthogroups that have one *Daucus carota* and three *Heracleum sosnowskyi* genes, left – whole range of variation of distances, 1-Mb bins, right – distances up to 1 Mb, 10-Kb bins. d: Distance between pairs of paralogues in orthogroups that have one *Daucus carota* and four *Heracleum sosnowskyi* genes, left – whole range of variation of distances, 1-Mb bins, right – distances up to 1 Mb, 10-Kb bins. e: Distance between pairs of paralogues in orthogroups that have one *Daucus carota* and five or more *Heracleum sosnowskyi* genes, left – whole range of variation of distances, 1-Mb bins, right – distances up to 1 Mb, 10-Kb bins.

**Table 3.** Genomic proximity of duplicated genes in *H. sosnowskyi*.

| All duplicated genes |  |  |  |  |  |
| --- | --- | --- | --- | --- | --- |
| Number of *H. sosnowskyi* genes in orthogroup | Number of orthogroups | Number of pairs of paralogs | Number of pairs of paralogs located on the same chromosome | Fraction of pairs of paralogs located on the same chromosome | Average distance between of pairs of paralogs located on the same chromosome |
| 2 | 1 225 | 1 225 | 449 | 0.366 | 4 192 923 |
| 3 | 244 | 732 | 318 | 0.434 | 3 008 666 |
| 4 | 115 | 690 | 365 | 0.529 | 6 715 887 |
| >=5 | 230 | 25 150 | 5 873 | 0.234 | 17 303 928 |
| Duplicated genes located on the same chromosome |  |  |  |  |  |
| Number of *H. sosnowskyi* genes in orthogroup | The fraction of paralogs located at a distance less than 10 Kb |  | The fraction of paralogs located at a distance less than 100 Kb |  | The fraction of paralogs located at a distance less than 1 Mb |
| 2 | 0.220 |  | 0.576 |  | 0.753 |
| 3 | 0.135 |  | 0.491 |  | 0.792 |
| 4 | 0.107 |  | 0.542 |  | 0.792 |
| >=5 | 0.059 |  | 0.254 |  | 0.519 |

The increase in the number of genes within a gene family creates the opportunity for the emergence of new functions (neofunctionalization) or diversification within the same function (subfunctionalization), which are the drivers of adaptation (Hanada *et al.*, 2008), (Zhao *et al.*, 2022). To determine whether the gene family expansions in hogweed are associated with some specific functions, we performed Gene Ontology enrichment analysis looking for the overrepresentation of certain GO categories in hogweed gene set compared to carrot gene set. We found 41 GO terms that were significantly overrepresented in hogweed. Most of these terms were associated with secondary metabolism (terpenoid biosynthesis, orcinol O-methyltransferase, and enolase reductase activity). Additionally, the proteasome system (ubiquitinated ligases, cullin) was affected by the expansion, as well as some families of transcription factors (Supplementary Table S6). For celery and coriander, there were also many GO terms that were overrepresented in their gene set compared to carrot (Supplementary Table S7, S8). Some GO terms were common for all three species, and some were unique, emphasizing that both shared and lineage-specific duplications contributed to the expansion of a gene set in the apioid superclade.

### Furanocoumarin biosynthesis pathway genes

A prominent feature of *H. sosnowskyi* is its ability to produce FCs, secondary metabolites, many of which have biological activities that range from adverse to beneficial. Furanocoumarin biosynthesis is found in several distant groups of plants, such as Rutaceae, Moraceae, Fabaceae and Apiaceae; it presumably independently emerged in each of them. Among the members of Apiaceae, the ability to synthesize FCs is confined almost exclusively to the apioid superclade, with a few exceptions. FC biosynthesis is a multistep process orchestrated by several enzymes. It starts from a general precursor, umbelliferone, which is then prenylated at the 6 or 8 position, resulting in a precursor of linear FC, demethylsuberosine, or angular FC, osthenol (Brown and Steck, 1973). They are converted, correspondingly, into marmesin and columbianethin. These two are the precursors of psoralen and angelicin, the backbone compounds for linear and angular FCs, respectively. Further modifications of those two basic structures (hydroxylation, methoxylation, prenylation, oxygenation, glycosylation, etc.) and their combinations create enormous FC diversity - up to 100 compounds in one species (Bourgaud *et al.*, 2014). Although the FC synthesis pathway has gained much attention, it has been studied in a piecemeal manner, with the enzymes corresponding to different steps being characterized using different model objects (*Ammi*, *Pastinaca*, *Peucedanum*, *Citrus*, and *Ficus)* (Munakata *et al.*, 2012; Hehmann *et al.*, 2004; Jian *et al.*, 2020; Larbat *et al.*, 2009; Villard *et al.*, 2021). For most of those objects, the genomic sequences are not available, and the identification of genes was based on transcriptomes or on screening using primers of conserved regions. We performed a genomewide search for the genes encoding the FC biosynthesis pathway and surveyed their expression in different aboveground organs of hogweed. The first step of FC synthesis is prenylation of umbelliferone, either in position 6 leading to the precursor of linear FC - demethylsuberosin or in position 8, leading to the precursor of angular FC - osthenol. These reactions are catalysed by dimethylallyldiphosphate: umbelliferone dimethylallyltransferase (UDT); earlier studies (Munakata *et al.*, 2016) have shown the existence of two UDTs catalysing 6- and 8-prenylation in *Pastinaca sativa*. They share strong sequence similarity (71% at the amino acid level) and are presumed to be the result of duplication. A single UDT capable of catalysing both reactions but with a strong preference towards C6-prenylation was found in parsley (Karamat *et al.*, 2014). In the hogweed genome, we found three genes with high similarity to U6DT/U8DT located on chromosome 6 in close proximity to one another; two of them - Hsosn_jg38723 and Hsosn_jg38725 - were very similar (99%), suggesting their origin by a recent tandem duplication. In coriander, there were two homologues of U6DT/U8DT, located on chromosome 5. This chromosome is largely syntenic to chromosome 6 of *H. sosnowskyi*. In celery, there were three genes located on chromosomes 9 and 11 and in an unplaced contig. Based on phylogenetic analysis, this diversity of UDT homologues in the apioid superclade is the result of a combination of shared and recent lineage-specific duplications (Supplementary Figure S13). In hogweed, three genes showed quite uniform expression across organs; two of them were expressed at a high level, while one of two U8DT homologues was expressed at a lower level than the other (Figure 4A). The next step of biosynthesis is the conversion of demethylsuberosin to marmesin for linear FC and of osthenol to columbianethin for angular FC. Marmesin synthase (MS) activity was shown biochemically in *Ammi majus* cell cultures (Hamerski and Matern, 1988). However, no corresponding enzyme was isolated in any species of Apiaceae. The gene encoding marmesin-synthase is cloned in *Ficus carica* (Moraceae). It belongs to the CYP76 family of the cytochrome P450 superfamily (Villard *et al.*, 2021). The hogweed gene with the highest similarity to marmesin synthase from *F. carica* is Hsosn_jg20931. It is, however, not expressed in leaves and inflorescence rays and expressed at very low levels in fruits (Figure 4A), while the content of psoralen, the compound formed on the next step after marmesin, is very high in these organs (Satzyperova and Komissarenko, 1978). It is thus unlikely that the protein encoded by Hsosn_jg20931 is MS. This gene belongs to a large orthogroup containing 18 hogweed genes, most of which (10) are the result of a recent tandem duplication (Supplementary Figure S14). Many of them were not expressed or expressed at very low levels. Four members of the orthogroup were expressed at high levels and are more plausible candidates for MS (Figure 4A). For columbianethin-synthase, although the conversion of osthenol to columbianetin in plant cells was shown (Brown and Steck, 1973), no corresponding enzyme was ever characterized. Based on the same logic as for U6DT/U8DT and psoralen/angelicin synthase, columbianethin-synthase should be looked for in the same group of genes to which marmesin-synthase belongs. The last stage of synthesis of core FCs - psoralen and angelicin - is catalysed by psoralen and angelicin synthase, respectively. These two enzymes are quite well characterized in Apiaceae - PS in *Ammi*, *Pastinaca*, *Apium*, and *Peucedanum* and AS in *Pastinaca* and *Peucedanum* (Larbat *et al.*, 2007; Larbat *et al.*, 2009; Jian *et al.*, 2020). In hogweed, the two genes with the highest similarity to PA and AS were two adjacent genes, Hsosn_jg70422 and Hsosn_jg70421, located on chromosome 2. This chromosome was largely colinear to chromosome 2 of celery and chromosome 8 of coriander (Supplementary Figure S11); congruent with this, these chromosomes carry two genes similar to PS and AS, also closely related to one another. In carrot, on chromosome 5, there were also two genes with high similarity to PS and AS (LOC108220230 and LOC108221492). However, phylogenetic analysis indicated that these two genes are a product of a recent tandem duplication rather than a more ancient duplication, giving rise to two genes in hogweed, celery and coriander. In carrot, there was also another gene with high similarity to psoralen synthase on chromosome 1 (Figure 4B). Based on the survey of expression levels, primary candidates for psoralen and angelicin synthase are the Hsosn_jg70422 and Hsosn_jg70421 genes because they were expressed at high levels (Figure 4A). The results of phylogenetic analysis suggested that Hsosn_jg70421 is PS and Hsosn_jg70422 is AS because they were grouped with corresponding genes from *Pastinaca sativa*, a close relative of hogweed (Figure 4B). Additional information can be drawn from the analysis of active sites. Dueholm and coworkers, in their study of the evolution of substrate recognition sites (SRSs) in cytochromes P450 from Apiaceae, identified key residues required for the enzyme to function as psoralen- or angelicin-synthase (Dueholm *et al.*, 2015). We analysed the protein sequences of Hsosn_jg70421 and Hsosn_jg70422 in the context of this study; the results also point to Hsosn_jg70422 as AS and Hsosn_jg70421 as PS (Supplementary Table S9). To experimentally test these predictions, we performed heterologous expression of the candidate genes in yeasts. *S. cerevisiae* strain 2805 was transformed by the vector containing the inserts of the complete coding sequence of Hsosn_jg70422 and Hsosn_jg70421 and the same plasmid with no insert. At 44 hours after the precursor addition, we analysed the samples obtained from the culture medium using LC-MS/MS to test for the presence of the expected products of enzymatic reactions. The mass spectrometric analysis showed that Hsosn_jg70421 is indeed a psoralen synthase; however, it did not confirm the angelicin synthase activity of Hsosn_jg70422 (Figure 4C, for detailed mass spectrometry results see Supplementary Figure S15).

**Figure 4.**
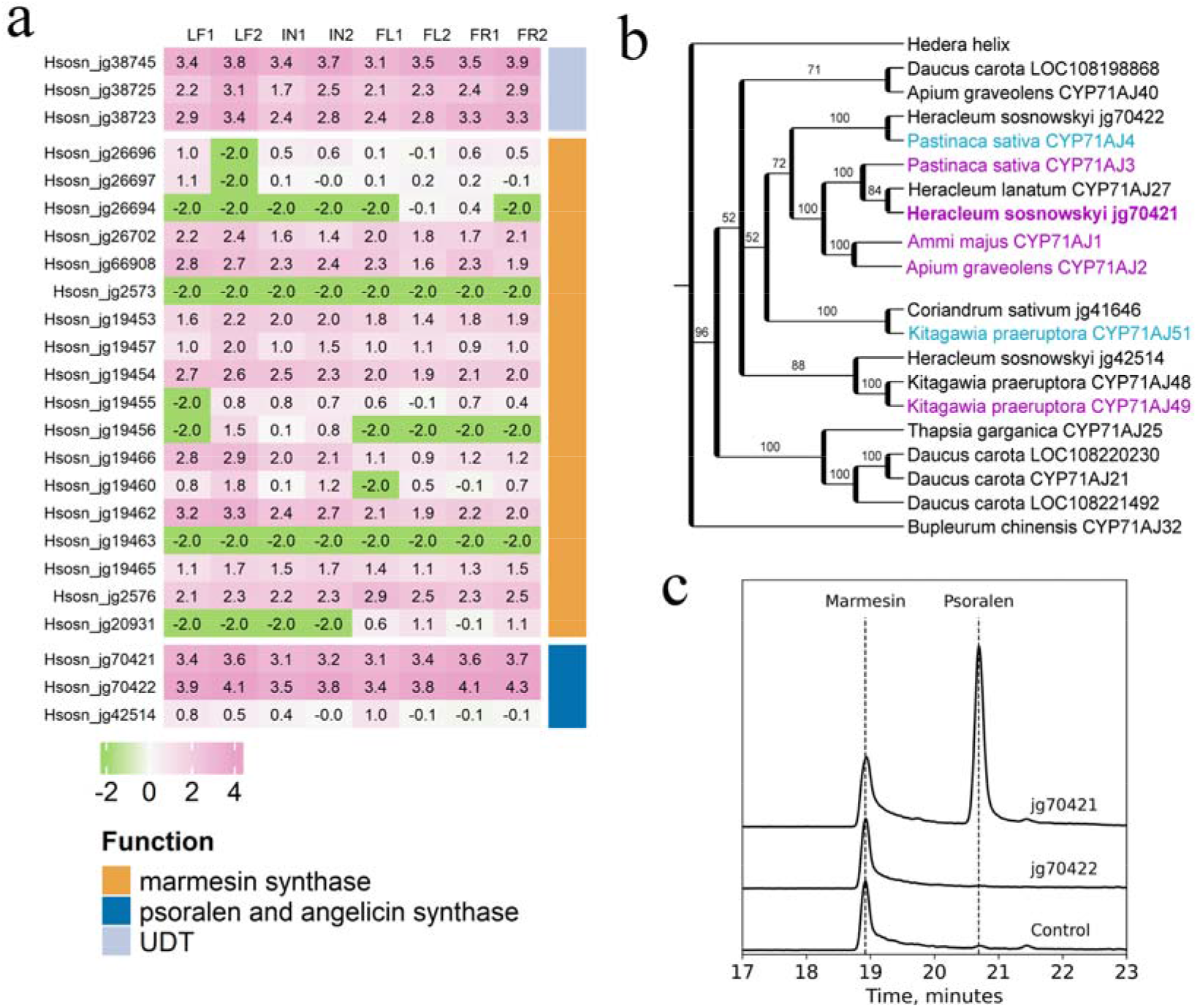
a: Heatmap showing the expression of furanocoumarin biosynthesis pathway homologues in *Heracleum sosnowskyi*. LF – leaf, IN – inflorescence rays, FL – flower, FR – fruits. Gene expression level was calculated as decimal logarithm of normalized expression level + pseudocount 0.01 (in order to avoid uncertainties when dealing with non-expressed genes). b: Phylogenetic tree of psoralen/angelicin-synthase-like genes in Apiaceae. The tree is based on maximum likelihood analysis of deduced amino acid sequences. Numbers above branches indicate bootstrap support (calculated based on 1000 replicates); nodes with support less than 50% are collapsed. The sequences for which function was shown experimentally are highlighted: purple for psoralen-synthase activity and cyan for angelicin-synthase activity. c: Mass spectrometry results for culture medium from transgenic yeasts expressing Hsosn_jg70421, Hsosn_jg70422 and the control strain transformed with empty vector.

## Methods

Detailed description of materials and methods is given in the the section “Supplemental experimental procedures”.

### Genome and transcriptome sequencing

DNA was extracted using the CTAB protocol (Doyle and Doyle, 1987) and sequenced on the Pacific Bioscience Sequel II platform (DNALink, South Korea). According to the HiC protocol, we first extracted nuclei from fresh leaves and then prepared libraries using the EpiTect Hi-C Kit (Qiagen, Netherlands). HiC libraries were sequenced on a NextSeq500 instrument (Illumina, USA) with a read length of 150+150 bp. For transcriptome sequencing, we extracted RNA from leaves, flowers, immature fruits and inflorescence rays using the RNEasy Mini kit (Qiagen, Netherlands). For Illumina library preparation, we used the NEBNextUltraII RNA library preparation kit (New England Biolabs, USA); libraries were sequenced on a HiSeq4000 instrument (Illumina, USA) with a read length of 150+150 bp. For nanoporesequencing, we first prepared double-stranded cDNA and then used it for sequencing on a MinION instrument with an LSK-109 kit (Oxford Nanopore Technologies, UK). Information on samples, their collection sites, read numbers and quality is available in Supplementary Table S1.

### Genome assembly and annotation

WGS and RNA-seq Illumina reads were trimmed by Fastp (Chen *et al.*, 2018). Oxford Nanopore reads were trimmed by Porechop (Wick, n.d.). The genome of *H. sosnowskyi* was assembled by Mabs-hifiasm (Schelkunov, 2022), Hifiasm (Cheng *et al.*, 2021), Flye (Kolmogorov *et al.*, 2019), Miniasm (Li, 2016), and Canu (Nurk *et al.*, 2020). After assembly, haplotypic duplication removal was performed by Purge_dups (Guan *et al.*, 2020). To compare the gene assembly quality in these assemblies, BUSCO (Simão *et al.*, 2015) and calculate_AG (Schelkunov, 2022) were used. The assembly with the most accurately assembled genes was the assembly made by Mabs without Purge_dups; this assembly was used in further analyses. Scaffolding using Hi-C reads was performed by Pin_hic (Guan *et al.*, 2021). Prior to annotation, repeats were masked in the scaffolds by RepeatModeler (Flynn *et al.*, 2020) and RepeatMasker (Smit *et al.*, n.d.). For annotation, we used BRAKER2 (Brůna *et al.*, 2021). To filter out false gene predictions made by BRAKER2, the following methods were used: 1) Proteins were aligned by DIAMOND (Buchfink *et al.*, 2021) to the NCBI nr database (Sayers *et al.*, 2021). Genes whose proteins had matches with e-values equal to or lower than 10^-5 were considered true genes. 2) Proteins were aligned by HMMER (Mistry et al., 2013) to the Pfam-A database (Mistry *et al.*, 2021). Genes whose proteins had matches with e-values equal to or lower than 10^-5 were considered true genes. 3) TPM was calculated for all genes by TPMCalculator (Vera Alvarez *et al.*, 2019). Then, the median TPM for genes whose proteins had matches in NCBI nr or Pfam was calculated. If a gene had a TPM above this median, it was considered a true gene even if it did not have matches in NCBI nr or Pfam. Thus, a gene predicted by BRAKER2 was considered a true gene if it was supported by homology or high expression. In addition, a gene was considered a false prediction and removed from the annotation if it met at least one of the following three criteria (even if it met some of the abovementioned criteria): 1) The protein encoded by a gene was shorter than 100 amino acids. 2) Most of the top 5 matches of the protein in NCBI nr belonged to taxa other than Embryophyta. 3) The gene belonged to a transposable element according to PANNZER2(Törönen and Holm, 2022). Genomes of *Coriandrum sativum*, *Apium graveolens* and *Daucus carota* were annotated with the same methods. Raw data, assemblies and annotations are available in the NCBI database under bioproject # PRJNA928505: https://www.ncbi.nlm.nih.gov/bioproject/?term=PRJNA928505

### Comparative genome analysis

Orthogroups were calculated for proteins of *H. sosnowskyi*, *C. sativum*, *A. graveolens*, *D. carota* and *Arabidopsis thaliana* by OrthoFinder (Emms and Kelly, 2019). The proteins of *A. thaliana* were obtained from TAIR10 (Berardini *et al.*, 2015). The distribution of the ratios of synonymous substitutions (“dS”) of paralogue pairs were calculated by WGD (Zwaenepoel and Van de Peer, 2019). Collinear blocks were found by MCScanX (Wang *et al.*, 2012). Collinear blocks were visualized by SynVisio (Bandi and Gutwin, Carl, 2020). The genomic proximity of duplicated genes was analysed using a set of custom scripts. The Gene Ontology (GO) terms enrichment analysis was made by GOATOOLS 1.3.1 (Klopfenstein *et al.*, 2018). The multiple hypothesis testing correction was performed using the method of Benjamini-Hochberg.

### Analysis of furanocoumarin pathway genes

As queries in the search for FC biosynthesis pathway genes, we used the sequences of the genes for which the function was shown experimentally, namely, dimethylallyldiphosphate: umbelliferone dimethylallyltransferases from *Pastinaca sativa* (accession numbers KM017083 and KM017084), marmesin synthase from *Ficus carica* (MW348922), psoralen-synthase from *Ammi majus* (AY532370) and angelicin-synthase from *Pastinaca sativa*(EF191021). For the genes of plants from Apiaceae, we used an 80% identity threshold and query coverage >90% for those that were outside 70% and 50%, respectively. Sequences were aligned using MUSCLE (Edgar, 2004); phylogenetic analysis was performed using IQ-tree (Minh *et al.*, 2020). Gene expression levels were normalized using median method (Anders and Huber, 2010) to decrease the influence of library size. Heatmap was pictured using R package “complexHeatmap” (Gu *et al.*, 2016).

### Expression of candidate genes in yeasts

We transformed yeast (S. *cerevisiae* strain 2805) with the expression vector pYeDP1/8-2 (Cullin and Pompon, 1988) carrying complete coding sequences of Hsosn_jg70421 and Hsosn_70422 amplified using *H. sosnoswkyi* leaf cDNA as a template. Expression of heterologous genes in *S. cerevisiae* cells was induced by galactose (Akiyoshi-Shibata *et al.*, 1991) and performed at 30°C for 18-44 hours. The efficiency of induction and the optimal time point for analysis were determined using qRT-PCR. The functional activity of the proteins of interest was tested using the addition of marmesin and columbianetin (precursors of psoralen and angelicin, respectively) to the culture medium during the cultivation of recombinant cells under conditions of induction of expression of heterologous genes. Detection of the expected compounds was performed using mass spectrometry of a culture medium at 44 hours after the addition of precursor on an Agilent 1290 Infinity LC system connected in series with an Agilent 6495 Triple Quadrupole mass spectrometer with an electrospray ionization source.

## Conclusions

The complexity of the plant genome, in particular, gene space, is well known to biologists. However, until recently, the study of its emergence and evolution was focused on one pattern – whole genome duplication and subsequent genome fractionation – while other patterns involving smaller-scale duplications remained mostly overlooked. This is mostly due to the technical challenges that prevent correct assembly and annotation, especially for recent duplications. The tremendous progress of sequencing technologies, in particular, the emergence of platforms capable of producing long reads, equips plant biologists with a tool capable of overcoming these challenges. Studies addressing the structure and function of tandem duplicated genes have begun to emerge (Liu *et al.*, 2023; Xu *et al.*, 2020), (Wang *et al.*, 2022); however, the contribution of tandem duplications to the total gene space is still unclear and probably differs between evolutionary lineages of flowering plants. Here, using a combination of long- and short-read sequencing, we assembled a chromosome-scale genome of Sosnowsky’s hogweed. We found that it has an unusually high number of genes and that this increase is mostly due to tandem duplications. Moreover, the high number of genes is shared between members of the apioid superclade - a large group within Apiaceae to which many medicinal and agricultural plants belong. We performed a genome-wide search of FC biosynthesis pathway genes and experimentally showed the function of one of the members of the CYP71 family genes as psoralen synthase. In addition to the expansion of our knowledge on the evolution of gene space in plants, this study opens an avenue for further genetic studies on Sosnowsky’s hogweed that are important for the biological control of this noxious invasive plant.

## Supporting information

Supplemental experimental procedures

## Data statement

Raw data were deposited into the NCBI SRA archive. Accession numbers are listed in Supplementary Table S1. Assembly and annotation are available under bioproject # PRJNA928505. Scripts and other associated data are available on Figshare: **https://figshare.com/articles/dataset/Genome_analysis_of_Heracleum_sosnowskyi_and_other_Apiaceae/21995738**.

## Acknowledgements

The study is supported by the Russian Science Foundation (sequencing, annotation and comparative genome analysis, project # 21-74-20145) and budgetary subsidies to IITP RAS (gene expression and phylogenetic analysis, project # FFNU-2022-0037) and MSU (experimental validation of protein function, theme “The study of intra- and intercellular interactions by molecular, cell biology, physiology, and mathematical methods and bioinformatics”) and the Ministry of Science and Higher Education of the Russian Federation (LC-MS/MS data analysis, agreement # 075-01645-22-06, project # 720000F.99.1.B385AV67000). The authors are grateful to Sergey Dudov (Lomonosov Moscow State University) for the photo of *H. sosnowskyi* and to American Journal Experts company for editing of the manuscript.

## Supplementary information

### Supplementary figures

Supplementary Figure S1.

Sina plots of sequencing coverage in exons of genes from single-copy and multicopy BUSCO orthogroups. Every dot is a gene. A peak in the multicopy diagrams that has half the coverage of the peak in the single-copy diagrams represents haplotypic duplications.

Supplementary Figure S2.

Map of the plastid genome of *H. sosnowskyi.* LSC – large single copy region, SSC – small single copy region, IRA and IRB – inverted repeat, regions A and B.

Supplementary Figure S3.

Phylogenetic tree of Apiaceae with a focus on tribe Tordylieae based on plastid genome sequences. Numbers above branches indicate bootstrap support (calculated based on 1000 replicates); nodes with support less than 50% are collapsed.

Supplementary Figure S4.

Long repeat content in the genomes of *H. sosnowskyi* and other Apiaceae. The diagram is based on the results of RepeatMasker.

Supplementary Figure S5.

*Heracleum sosnowskyi* genome map showing the density of repeats and protein-coding genes. Supplementary Figure S6.

*Coriandrum sativum*, genome map showing the density of repeats and protein-coding genes Supplementary Figure S7

*Apium graveolens* genome map showing the density of repeats and protein-coding genes Supplementary Figure S8

*Daucus carota* genome map showing the density of repeats and protein-coding genes Supplementary Figure S9

Protein-coding genes in *H. sosnowskyi* annotation, supported by homology and RNA-seq data.

Supplementary Figure S10

Dependence between gene copy number per orthogroup and sequencing coverage in genes of *Heracleum sosnowskyi.*“Sequencing coverage” of a gene is the median coverage in the gene’s exons by genomic reads. Box borders denote the 25th and 75th percentiles, whiskers denote the 5th and 95th percentiles, and the middle line denotes the median.

Supplementary Figure S11.

Collinear blocks between genomes of *Daucus carota* (chromosomes denoted as dc1 – dc9) and members of apioid superclade: *Apium graveolens* (chromosomes denoted as ag1 – ag9) and *Coriandrum sativum* (chromosomes denoted as cs1 – cs11). Only blocks that contain at least 20 pairs of homologous genes are shown. Coordinates are indicated in Mb.

Supplementary Figure S12

Genomic proximity of duplicated genes in *Apium graveolens* and *Coriandrum sativum*.

a: Distance between pairs of paralogues in orthogroups that have one *Daucus carota* and two *Apium graveolens/Coriandrum sativum* genes, left – whole range of variation of distances, 1-Mb bins, right – distances up to 1 Mb, 10-Kb bins.

b: Distance between pairs of paralogues in orthogroups that have one *Daucus carota* and three *Apium graveolens*/*Coriandrum sativum* genes, left – whole range of variation of distances, 1-Mb bins, right – distances up to 1 Mb, 10-Kb bins.

c: Distance between pairs of paralogues in orthogroups that have one *Daucus carota* and four *Apium graveolens/Coriandrum sativum* genes, left – whole range of variation of distances, 1-Mb bins, right – distances up to 1 Mb, 10-Kb bins.

d: Distance between pairs of paralogues in orthogroups that have one *Daucus carota* and five or more *Apium graveolens*/*Coriandrum sativum* genes, left – whole range of variation of distances, 1-Mb bins, right – distances up to 1 Mb, 10-Kb bins.

Supplementary Figure S13.

Phylogenetic tree of UDT homologues in Apiaceae. Numbers above branches indicate bootstrap support (calculated based on 1000 replicates); nodes with support less than 50% are collapsed.

Supplementary Figure S14.

Phylogenetic tree of marmesin-synthase homologues in Apiaceae. Numbers above branches indicate bootstrap support (calculated based on 1000 replicates); nodes with support less than 50% are collapsed.

Supplementary Figure S15.

Mass spectrometry results for transgenic yeasts expressing jg70421 and jg70422 and the control strain.

### Supplementary Tables

Supplementary Table S1. Information on samples, their collection sites and sequencing data.

Supplementary Table S2. Information on *Heracleum sosnowskyi* contigs produced by various genome assemblers. BUSCO statistics correspond to the “eudicots” dataset of BUSCO.

Supplementary Table S3. Analysis of random pairs: the probability of being on the same chromosome and distance between pairs where both genes are on the same chromosome. A total of 10 000 random pairs are analysed.

Supplementary Table S4. Genomic proximity of duplicated genes in coriander and celery. Supplementary Table S5. Genomic proximity of duplicated genes located on the same chromosome in coriander and celery.

Supplementary Table S6. GO enrichment for the gene set of *Heracleum sosnowskyi*compared with *Daucus carota*.

Supplementary Table S7. GO enrichment the gene set of *Apium graveolens* compared with *Daucus carota*.

Supplementary Table S8. GO enrichment the gene set of *Coriandrum sativum* compared with *Daucus carota*.

Supplementary Table S9. Psoralen and angelicin-synthase residues in substrate recognition sites (SRS) of candidate proteins compared to proteins for which the function was shown experimentally.

