## Supplementary Table 1 for "The genome of the toxic invasive species *Heracleum sosnowskyi* carries an increased number of genes despite absence of recent whole-genome duplications"

Supplementary table 1. Information on samples, their collection sites and sequencing data

| **Sample ID** | **Sample type** | **Organ** | **Collection site** | **Sequencing platform** | **Number of reads** | **Mean quality (only for Illumina)** | **Sequencing Read Archive accession code** |
| --- | --- | --- | --- | --- | --- | --- | --- |
| Hsosn_3 | genome (WGS) | leaf | 55°38'23.22"N  36°59'54.92"E | PacificBioscience, Sequel II | 1382 306 |  | SRR23251371,  SRR23251372 |
| Hsosn_3 | genome (HiC) | leaf | 55°38'23.22"N  36°59'54.92"E | Illumina, Nextseq500 | 372 768 859 | 34.4 | SRR23251383,  SRR23251384 |
| Hsosn_1_fruit | RNA | fruit | 62°50'46.19"N  34°46'14.51"E | Illumina, Hiseq4000 | 27 430 879 | 38.1 | SRR23251370 |
|  |  |  |  | Oxford Nanopore technologies, Minion | 80 308 |  | SRR23251369 |
| Hsosn_2_leaf | RNA | leaf | 62°50'46.19"N  34°46'14.51"E | Illumina, Hiseq4000 | 18 973 391 | 37.8 | SRR23251368 |
|  |  |  |  | Oxford Nanopore technologies, Minion | 58 621 |  | SRR23251367 |
| Hsosn_3_flower | RNA | flower | 62°50'46.19"N  34°46'14.51"E | Illumina, Hiseq4000 | 17 346 034 | 37.6 | SRR23251366 |
|  |  |  |  | Oxford Nanopore technologies, Minion | 85 732 |  | SRR23251365 |
| Hsosn_4_inflor | RNA | inflorescence rays | 62°50'46.19"N  34°46'14.51"E | Illumina, Hiseq4000 | 17 728 772 | 37.7 | SRR23251382 |
|  |  |  |  | Oxford Nanopore technologies, Minion | 214 817 |  | SRR23251381 |
| Hsosn_9_fruit | RNA | fruit | 55°38'23.22"N  36°59'54.92"E | Illumina, Hiseq4000 | 20 520 070 | 37.9 | SRR23251380 |
|  |  |  |  | Oxford Nanopore technologies, Minion | 107 664 |  | SRR23251379 |
| Hsosn_10_leaf | RNA | leaf | 55°38'23.22"N  36°59'54.92"E | Illumina, Hiseq4000 | 17 563 646 | 37.8 | SRR23251378 |
|  |  |  |  | Oxford Nanopore technologies, Minion | 147 623 |  | SRR23251377 |
| Hsosn_11_flower | RNA | flower | 55°38'23.22"N  36°59'54.92"E | Illumina, Hiseq4000 | 20 893 784 | 37.8 | SRR23251376 |
|  |  |  |  | Oxford Nanopore technologies, Minion | 118 280 |  | SRR23251375 |
| Hsosn_12_inflor | RNA | inflorescencerays | 55°38'23.22"N  36°59'54.92"E | Illumina, Hiseq4000 | 21 096 069 | 37.7 | SRR23251374 |
|  |  |  |  | Oxford Nanopore technologies, Minion | 4 337 |  | SRR23251373 |
