## Supplementary Table 2 for "The genome of the toxic invasive species *Heracleum sosnowskyi* carries an increased number of genes despite absence of recent whole-genome duplications"

Supplementary Table 2. Information on *Heracleum sosnowskyi* contigs produced by various genome assemblers. BUSCO statistics correspond to the "eudicots" dataset of BUSCO.

† Calculated by the program calculate_AG from a suite of programs Mabs. "True multicopy orthogroups" and "False multicopy orthogroups" are distinguished based on their coverage. "True multicopy orthogroups" are composed of genes that are paralogs, while "false multicopy orthogroups" are composed of genes that are erroneous duplications that are mistakes of a genome assembler. The criteria for "completeness" are the same as in BUSCO, i.e. a gene is considered assembled completely if it passes orthogroup-specific thresholds on bit score and length.

| Assembler | N50 (bp) | Sum of contigs' lengths (bp) | BUSCO results | Number of completely assembled BUSCO genes | | |
| --- | --- | --- | --- | --- | --- | --- |
|  |  |  |  | single-copy † | in true multicopy orthogroups† | in false multicopy orthogroups† |
| Mabs-hifiasm | 22 773 521 | 1 630 587 584 | C:98.3%[S:89.8%,D:8.5%], F:0.3%,M:1.4%,n:2326 | 1 706 | 845 | 116 |
| Hifiasm | 14 169 039 | 1 807 160 894 | C:98.5%[S:81.5%,D:17.0%], F:0.3%,M:1.2%,n:2326 | 1 642 | 812 | 563 |
| Flye | 901 982 | 2 641 773 227 | C:98.6%[S:27.7%,D:70.9%], F:0.3%,M:1.1%,n:2326 | 487 | 338 | 3 753 |
| Miniasm | 63 703 | 2 754 827 650 | C:96.6%[S:52.1%,D:44.5%], F:0.7%,M:2.7%,n:2326 | 965 | 497 | 2 605 |
| Canu | 1 941 778 | 2 869 254 115 | C:98.6%[S:22.9%,D:75.7%], F:0.3%,M:1.1%,n:2326 | 406 | 333 | 3 942 |
| Mabs-hifiasm+Purge_dups | 53 959 280 | 178 745 153 | C:11.4%[S:11.0%,D:0.4%], F:0.6%,M:88.0%,n:2326 | 201 | 34 | 2 |
| Hifiasm+Purge_dups | 14 212 445 | 469 846 757 | C:36.4%[S:34.3%,D:2.1%], F:0.9%,M:62.7%,n:2326 | 659 | 191 | 17 |
| Flye + Purge_dups | 1 190 311 | 1 472 349 734 | C:94.7%[S:84.1%,D:10.6%], F:0.6%,M:4.7%,n:2326 | 1 580 | 776 | 246 |
| Miniasm + Purge_dups | 77 136 | 1 699 258 648 | C:89.5%[S:70.3%,D:19.2%],  F:1.5%,M:9.0%,n:2326 | 1 317 | 660 | 835 |
| Canu + Purge_dups | 2 425 164 | 969 027 090 | C:76.8%[S:70.2%,D:6.6%],  F:1.1%,M:22.1%,n:2326 | 1 355 | 551 | 89 |
