## Supplementary Table 3 for "The genome of the toxic invasive species *Heracleum sosnowskyi* carries an increased number of genes despite absence of recent whole-genome duplications"

Supplementary table 3. Analysis of random pairs: the probability of being on the same chromosome and distance between pairs where both genes are on the same chromosome. 10 000 random pairs are analyzed.

| Species | The fraction of random pairs located on the same chromosome | Average distance between random pairs located on the same chromosome, bp |
| --- | --- | --- |
| Heracleum sosnowskyi | 0.090 | 49 556 824 |
| Coriandrum sativum | 0.100 | 57 746 439 |
| Apium graveolens | 0.091 | 111 293 311 |
