## Supplementary Table 4 for "The genome of the toxic invasive species *Heracleum sosnowskyi* carries an increased number of genes despite absence of recent whole-genome duplications"

Supplementary table 4. Genomic proximity of duplicated genes in coriander and celery

a) *Coriandrum sativum*

| Number of *Coriandrum sativum* genes in orthogroup | Number of orthogroups | Number of pairs of paralogs | Number of pairs of paralogs located on the same chromosome | Fraction of pairs of paralogs located on the same chromosome | Average distance between of pairs of paralogs located on the same chromosome |
| --- | --- | --- | --- | --- | --- |
| 2 | 1 306 | 1 306 | 714 | 0.547 | 2 578 101 |
| 3 | 197 | 591 | 301 | 0.509 | 5 107 755 |
| 4 | 59 | 354 | 182 | 0.514 | 1 571 784 |
| >=5 | 117 | 50 632 | 6 226 | 0.123 | 37 511 744 |

b) *Apium graveolens*

| Number of *Apium graveolens* genes in orthogroup | Number of orthogroups | Number of pairs of paralogs | Number of pairs of paralogs located on the same chromosome | Fraction of pairs of paralogs located on the same chromosome | Average distance between of pairs of paralogs located on the same chromosome |
| --- | --- | --- | --- | --- | --- |
| 2 | 982 | 982 | 337 | 0.343 | 10 007 432 |
| 3 | 182 | 546 | 249 | 0.456 | 12 989 717 |
| 4 | 86 | 516 | 201 | 0.390 | 17 079 734 |
| >=5 | 181 | 61 473 | 9 385 | 0.153 | 54 127 083 |
