## Supplementary Table 5 for "The genome of the toxic invasive species *Heracleum sosnowskyi* carries an increased number of genes despite absence of recent whole-genome duplications"

a) *Coriandrum sativum*

| Number of *Coriandrum sativum* genes in orthogroup | The fraction of paralogs located at a distance less than 10 Kb | The fraction of paralogs located at a distance less than 100 Kb | The fraction of paralogs located at a distance less than 1 Mb |
| --- | --- | --- | --- |
| 2 | 0.239 | 0.618 | 0.909 |
| 3 | 0.352 | 0.674 | 0.874 |
| 4 | 0.220 | 0.621 | 0.945 |
| >=5 | 0.049 | 0.138 | 0.244 |

b) *Apium graveolens*

| Number of *Apium graveolens* genes in orthogroup | The fraction of paralogs located at a distance less than 10 Kb | The fraction of paralogs located at a distance less than 100 Kb | The fraction of paralogs located at a distance less than 1 Mb |
| --- | --- | --- | --- |
| 2 | 0.291 | 0.608 | 0.819 |
| 3 | 0.213 | 0.538 | 0.827 |
| 4 | 0.159 | 0.662 | 0.338 |
| >=5 | 0.030 | 0.139 | 0.317 |

Supplementary table 5. Genomic proximity of duplicated genes located on the same chromosome, in coriander and celery.
