## Supplementary Table 9 for "The genome of the toxic invasive species *Heracleum sosnowskyi* carries an increased number of genes despite absence of recent whole-genome duplications"

| Amino acid | *jg70421* | *jg70422* | Psoralen-synthase | | | | Angelicin-synthase | |
| --- | --- | --- | --- | --- | --- | --- | --- | --- |
|  |  |  | *Ammi majus* (CYP71AJ1)^1^ | *Peucedanum praeruptorum* (CYP71AJ49)^2^ | *Apium graveolens* (CYP71AJ2)^3^ | *Pastinaca sativa* (CYP71AJ3)^3^ | *Peucedanum praeruptorum* (CYP71AJ51)^2^ | *Pastinaca sativa* (CYP71AJ4)^3^ |
| Substrate recognition site 1 | | | | | | | | |
| Arg104 | R | R | R | R | R | R | R | R |
| Val121 | V | A | V | A | V | V | G | A |
| Met120 | M | V | M | V | M | M | V | V |
| Substrate recognition site 4 | | | | | | | | |
| Ala297 | A | G | A | G | A | A | A | G |
| Thr301 | T | T | T | T | T | T | T | T |
| 297-303 (A/G)GX(D/E)T(T/S) | AGTETT | GGTETT | AGTETI | GGTETT | AGTETI | AGTETI | AGTETT | GGTETT |
| Substrate recognition site 5 | | | | | | | | |
| Thr361 | T | T | T | I | T | T | T | T |
| 359-361 YFT | YFT | YIT | YFT | YFI | YFT | YFT | YIT | YIT |
| Ala362 | A | A | A | A | A | A | A | A |
| Leu365 | L | L | L | L | L | L | L | L |
| Val366 | V | L | V | S | V | V | V | L |
| Pro367 | P | P | P | P | P | P | P | P |
| Substrate recognition site 6 | | | | | | | | |
| Thr479 | T | S | T | T | T | T | S | S |

Supplementary table 9. Key psoralen and angelicin-synthase residues in substrate recognition sites (SRS) of candidate proteins compared to proteins for which the function was shown experimentally

References.

^1^**Larbat, R., Kellner, S., Specker, S., Hehn, A., Gontier, E., Hans, J., Bourgaud, F. and Matern, U.** (2007) Molecular cloning and functional characterization of psoralen synthase, the first committed monooxygenase of furanocoumarin biosynthesis. *J. Biol. Chem.*, **282**, 542–554.

^2^**Jian, X., Zhao, Y., Wang, Z., Li, S., Li, L., Luo, J. and Kong, L.** (2020) Two CYP71AJ enzymes function as psoralen synthase and angelicin synthase in the biosynthesis of furanocoumarins in *Peucedanum praeruptorum* Dunn. *Plant Mol. Biol.*, **104**, 327–337.(note that *Peucedanum praeruptorum* is now known under the name of *Kitagawia praeruptora*).

^3^**Larbat, R., Hehn, A., Hans, J., Schneider, S., Jugdé, H., Schneider, B., Matern, U. and Bourgaud, F.** (2009) Isolation and functional characterization of *CYP71AJ4* encoding for the first P450 monooxygenase of angular furanocoumarin biosynthesis. *J. Biol. Chem.*, **284**, 4776–4785.
