## Supplementary Figures for "The genome of the toxic invasive species *Heracleum sosnowskyi* carries an increased number of genes despite absence of recent whole-genome duplications"

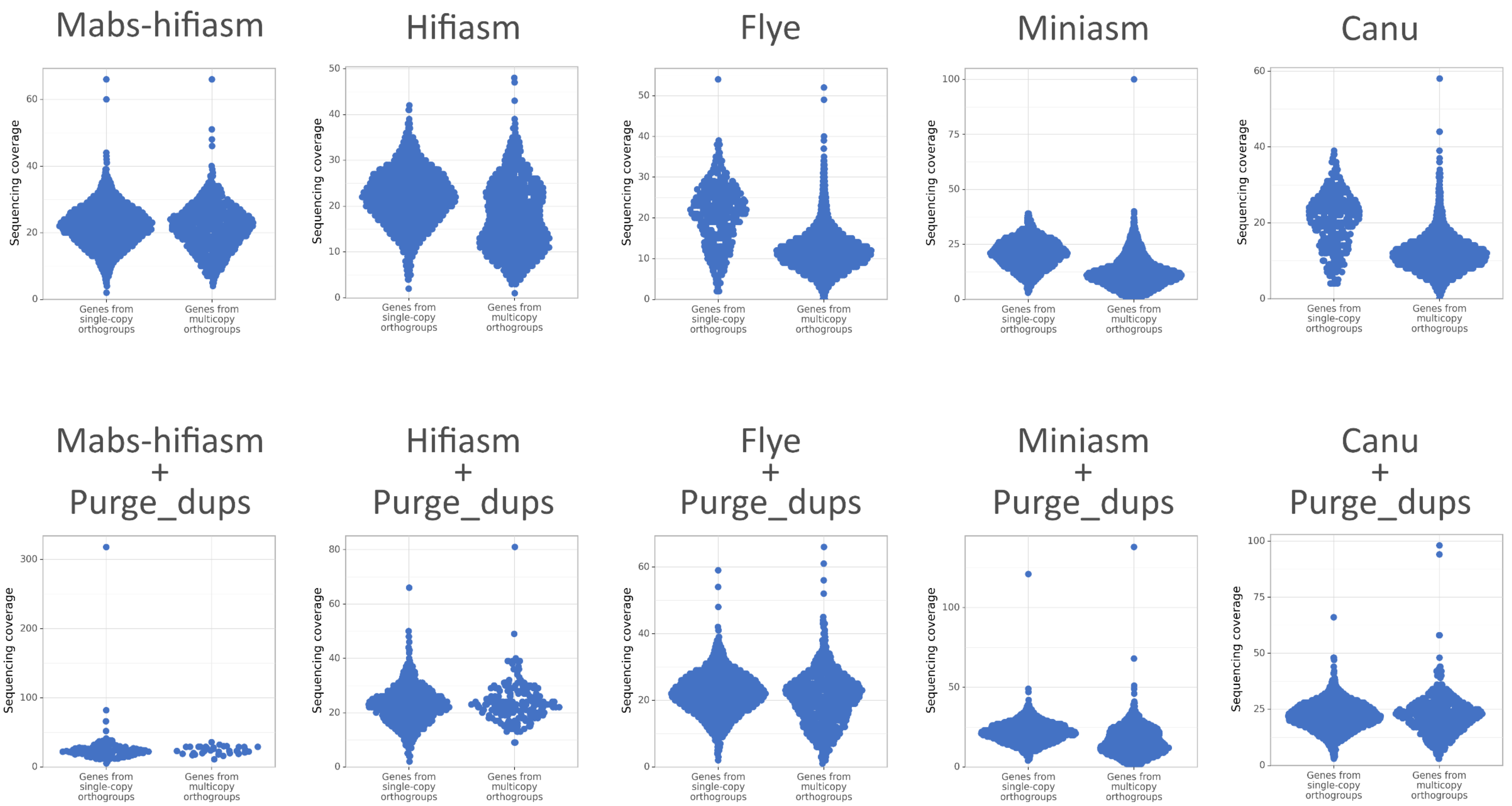


**Supplementary Figure 1.** Sinaplots of sequencing coverage in exons of genes from single-copy and multicopy BUSCO orthogroups. Every dot is a gene. A peak in the multicopy diagrams that has half the coverage of the peak in the single-copy diagrams represents haplotypic duplications.

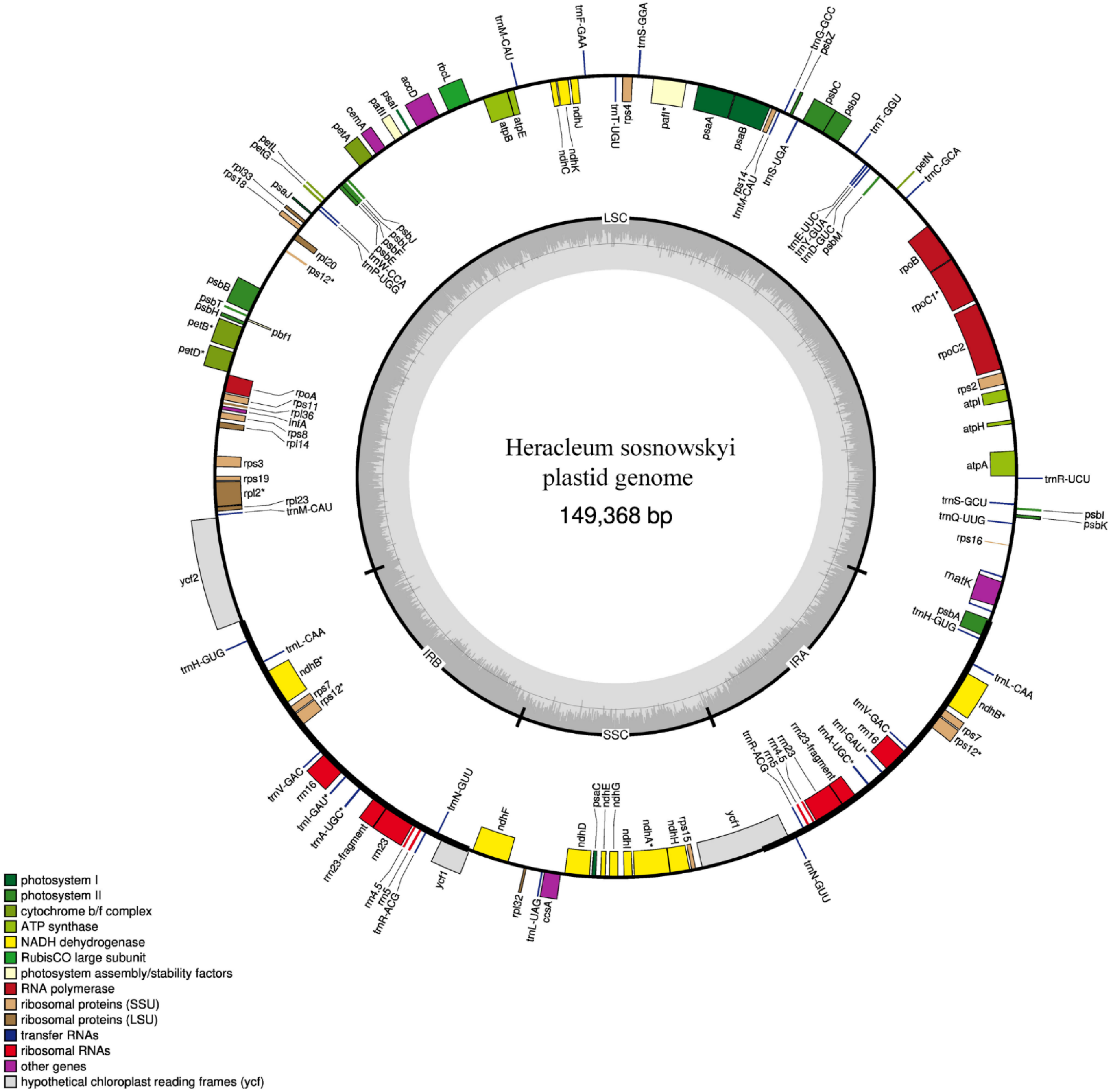


**Supplementary Figure 2.** Map of the plastid genome of *H. sosnowskyi*. LSC – large single copy region, SSC – small single copy region, IRA and IRB – inverted repeat, regions A and B.

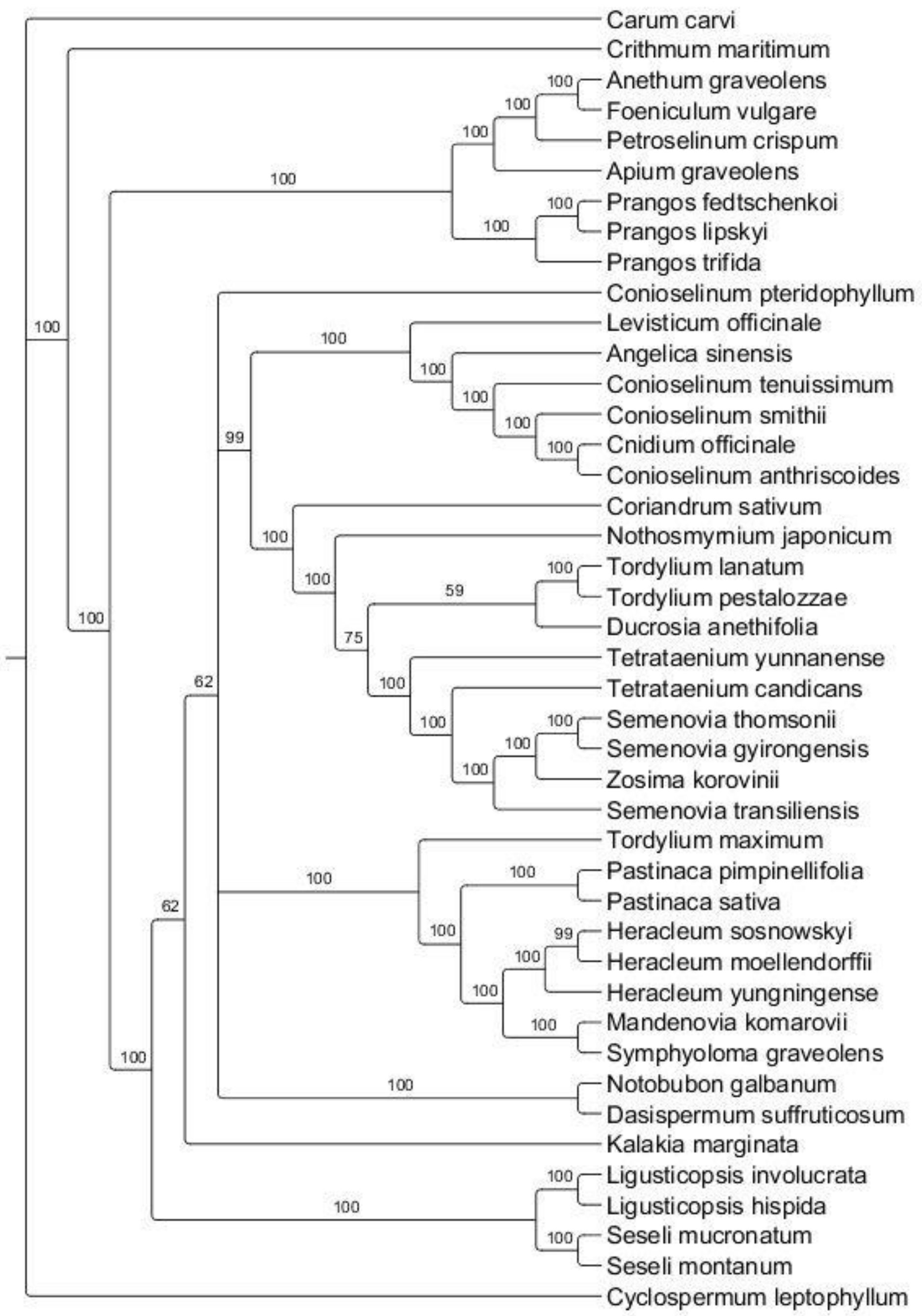


**Supplementary Figure 3.** Phylogenetic tree of Apiaceae with focus on tribe Tordylieae based on plastid genome sequences.

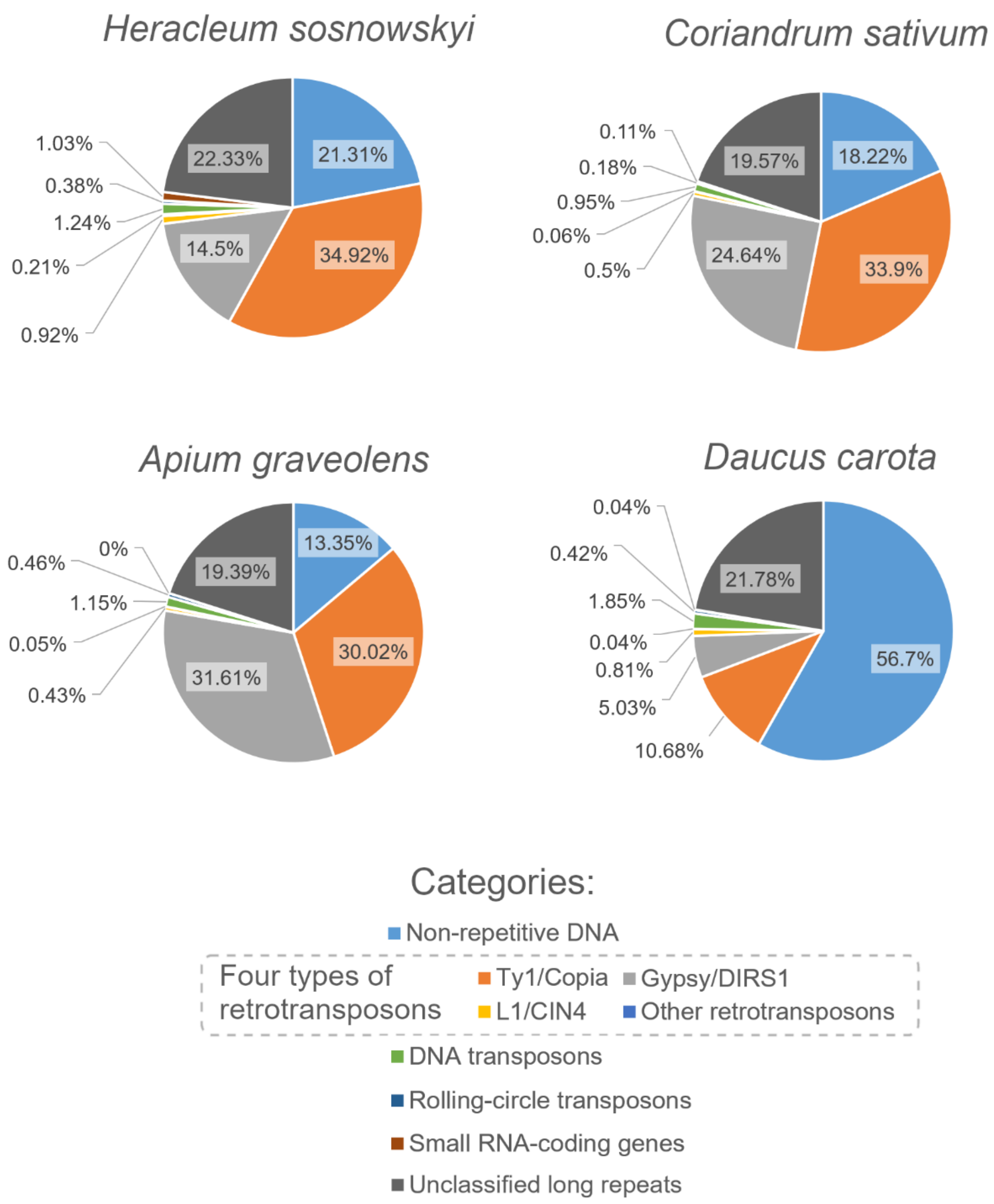


**Supplementary Figure 4.** Long repeat content in the genomes of *H. sosnowskyi* and other Apiaceae. Diagram is based on the results of RepeatMasker.

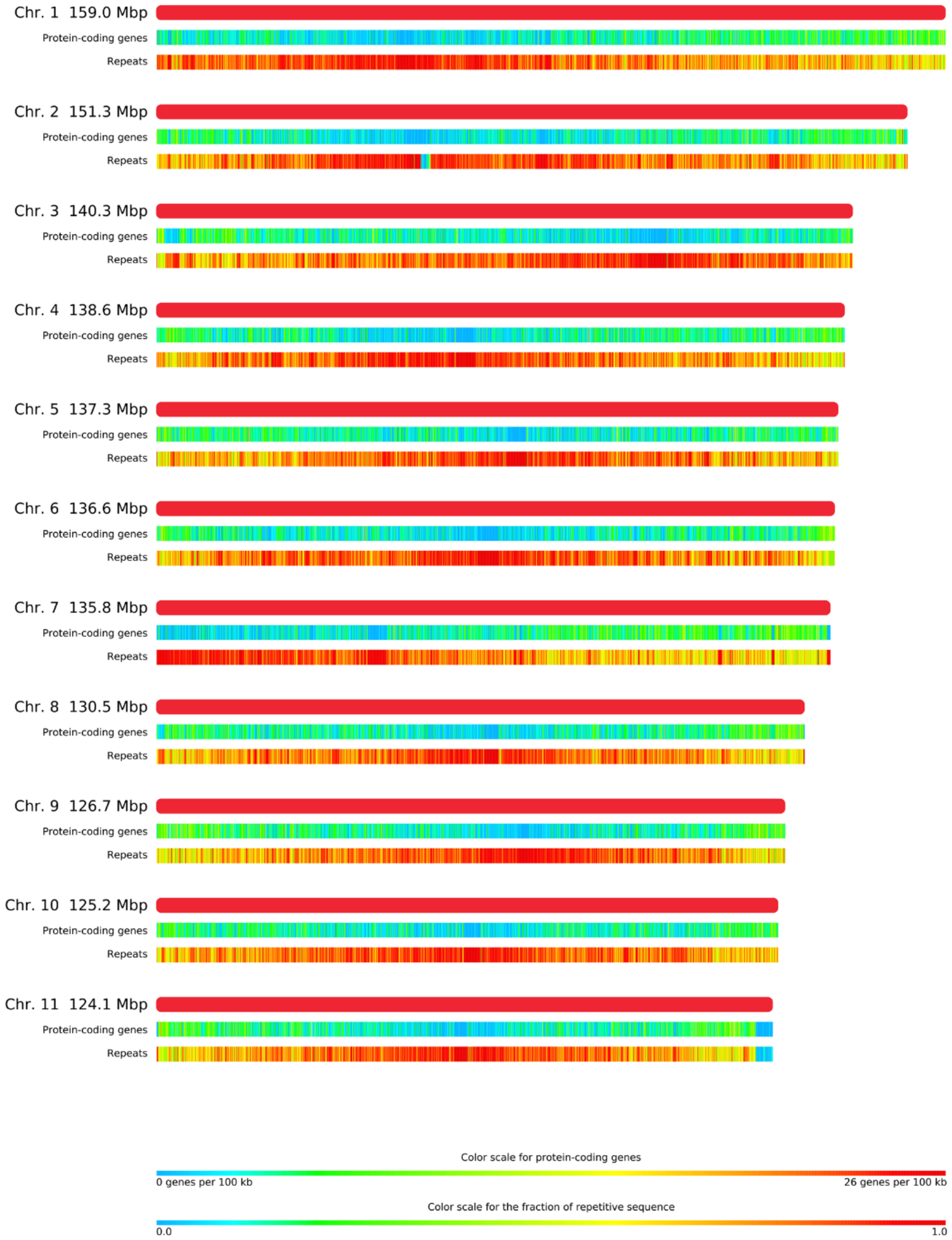


**Supplementary Figure 5.** *Heracleum sosnowskyi*, genome map showing the density of repeats and protein-coding genes.

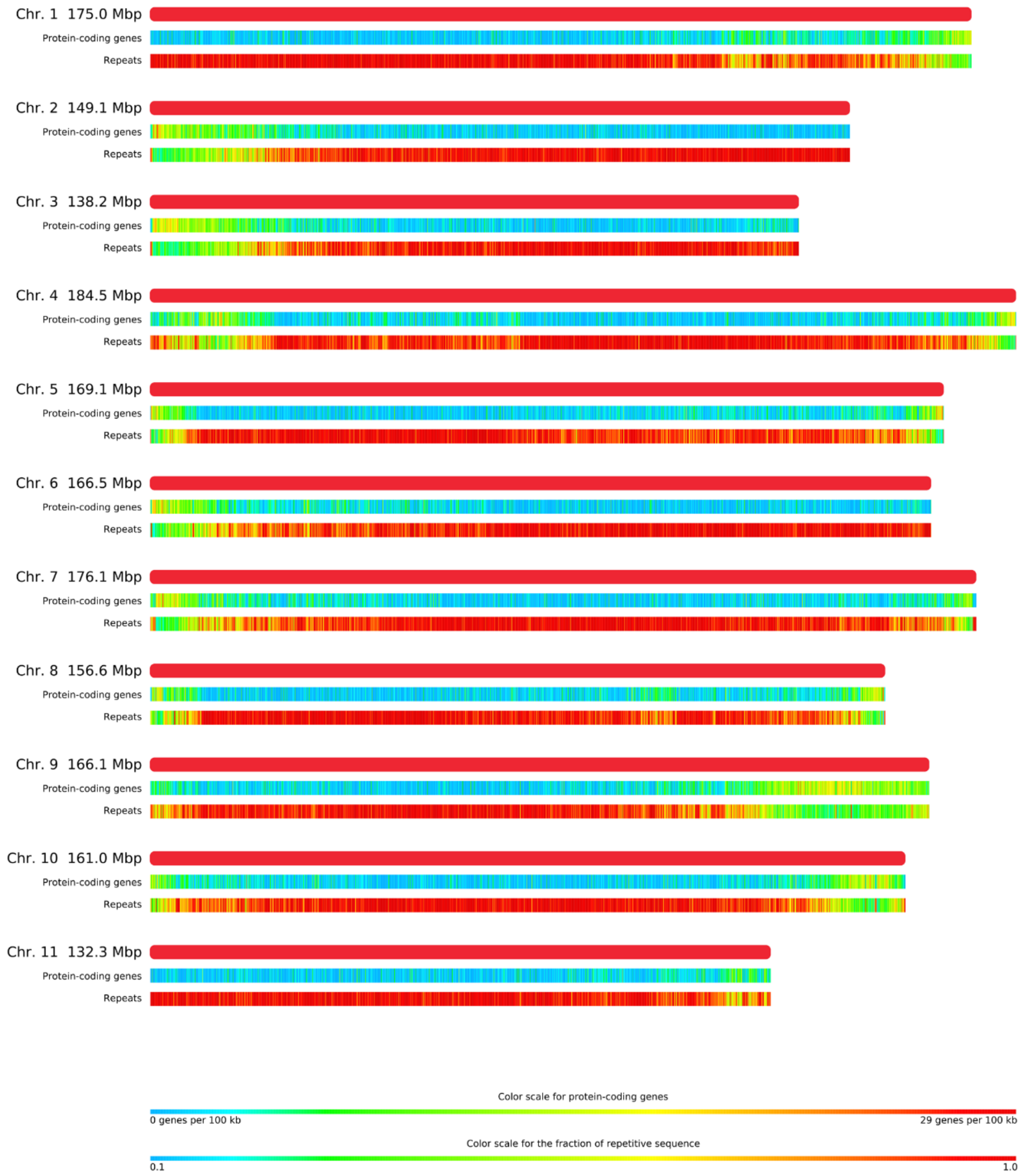


**Supplementary Figure 6.** *Coriandrum sativum*, genome map showing the density of repeats and protein-coding genes

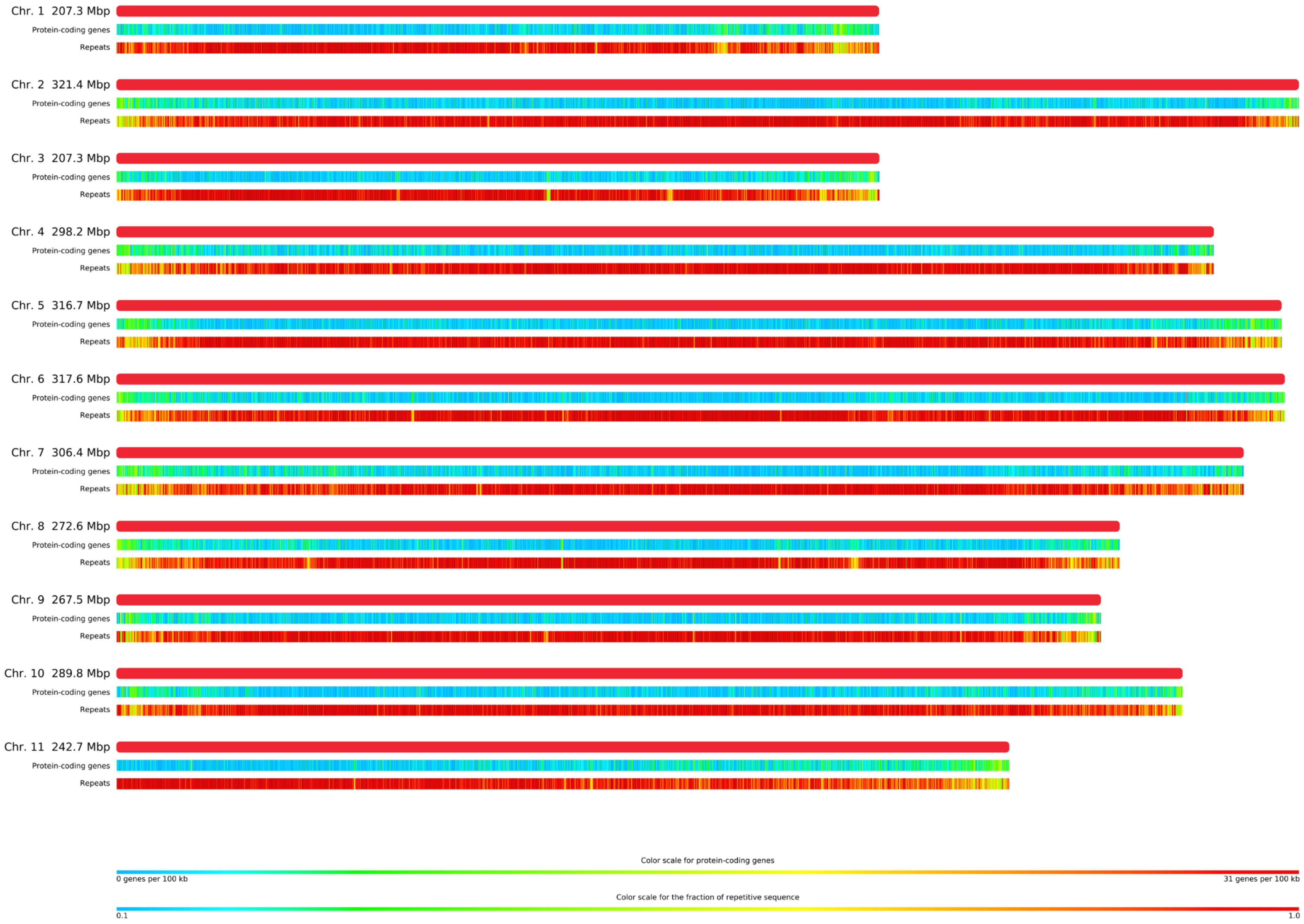


**Supplementary Figure 7.** *Apium graveolens*, genome map showing the density of repeats and protein-coding genes

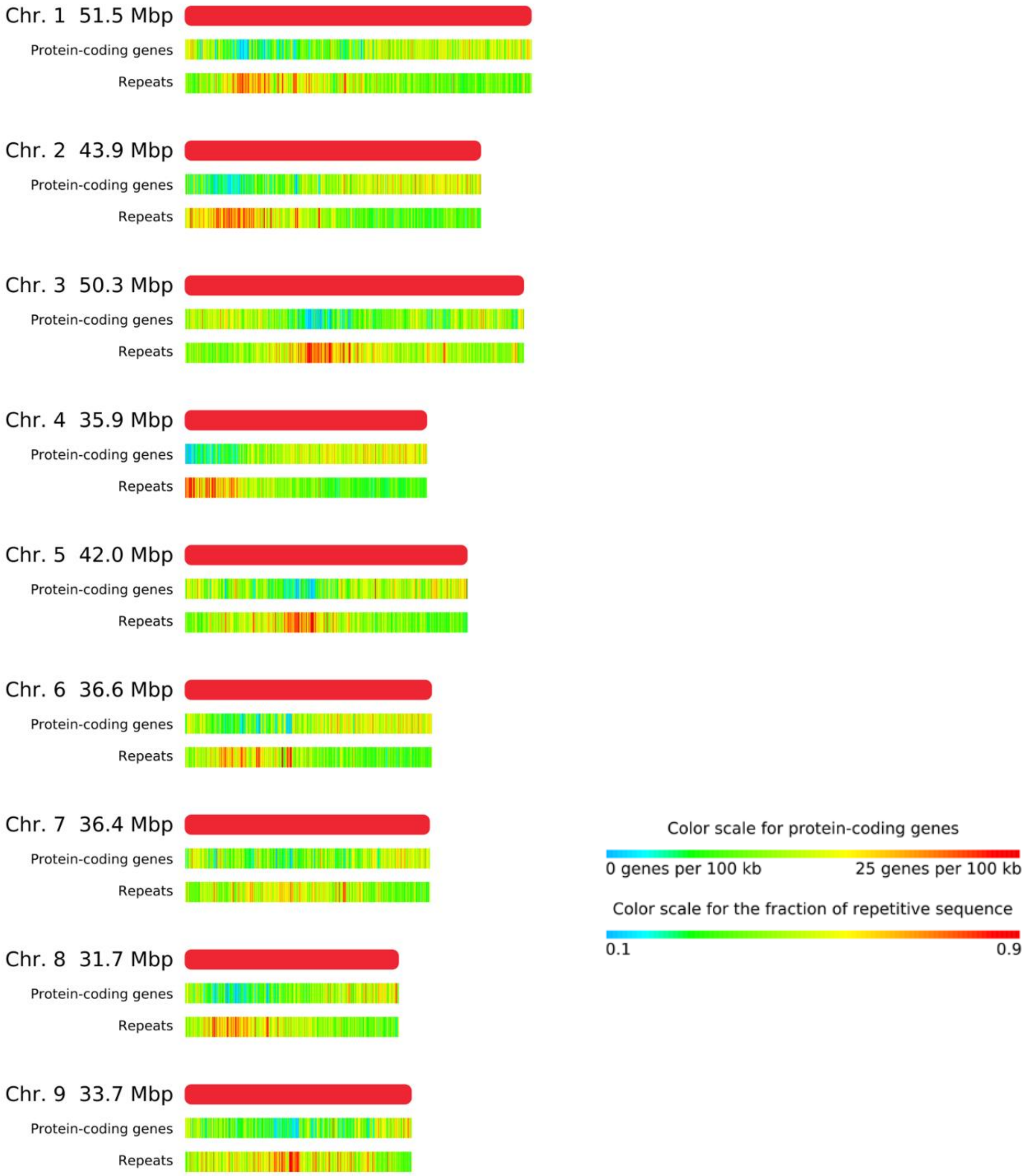


**Supplementary Figure 8.** *Daucus carota*, genome map showing the density of repeats and protein-coding genes

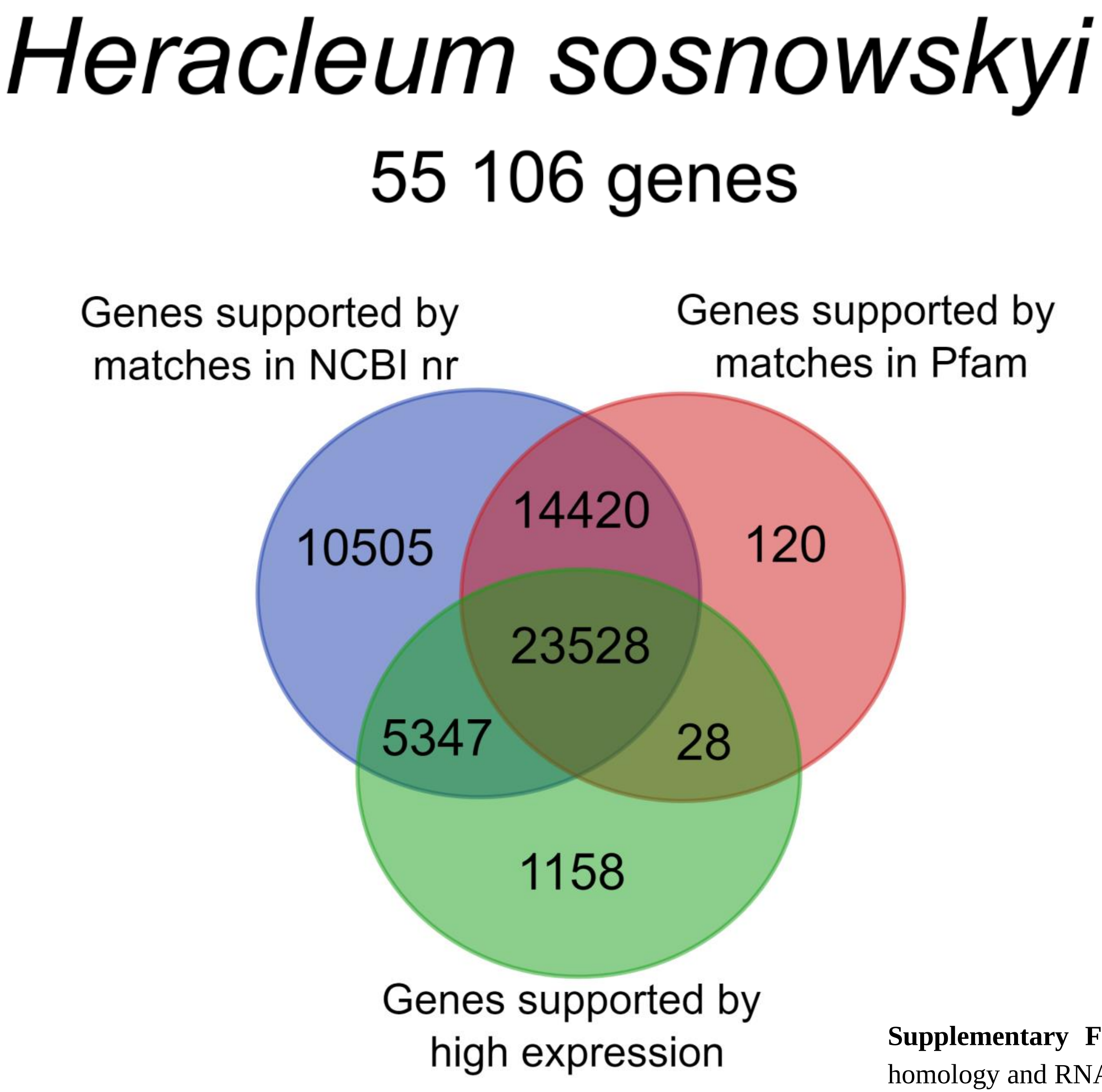


**Supplementary Figure 9.** Protein-coding genes in *H. sosnowskyi* annotation, support by homology and RNA-seq data.

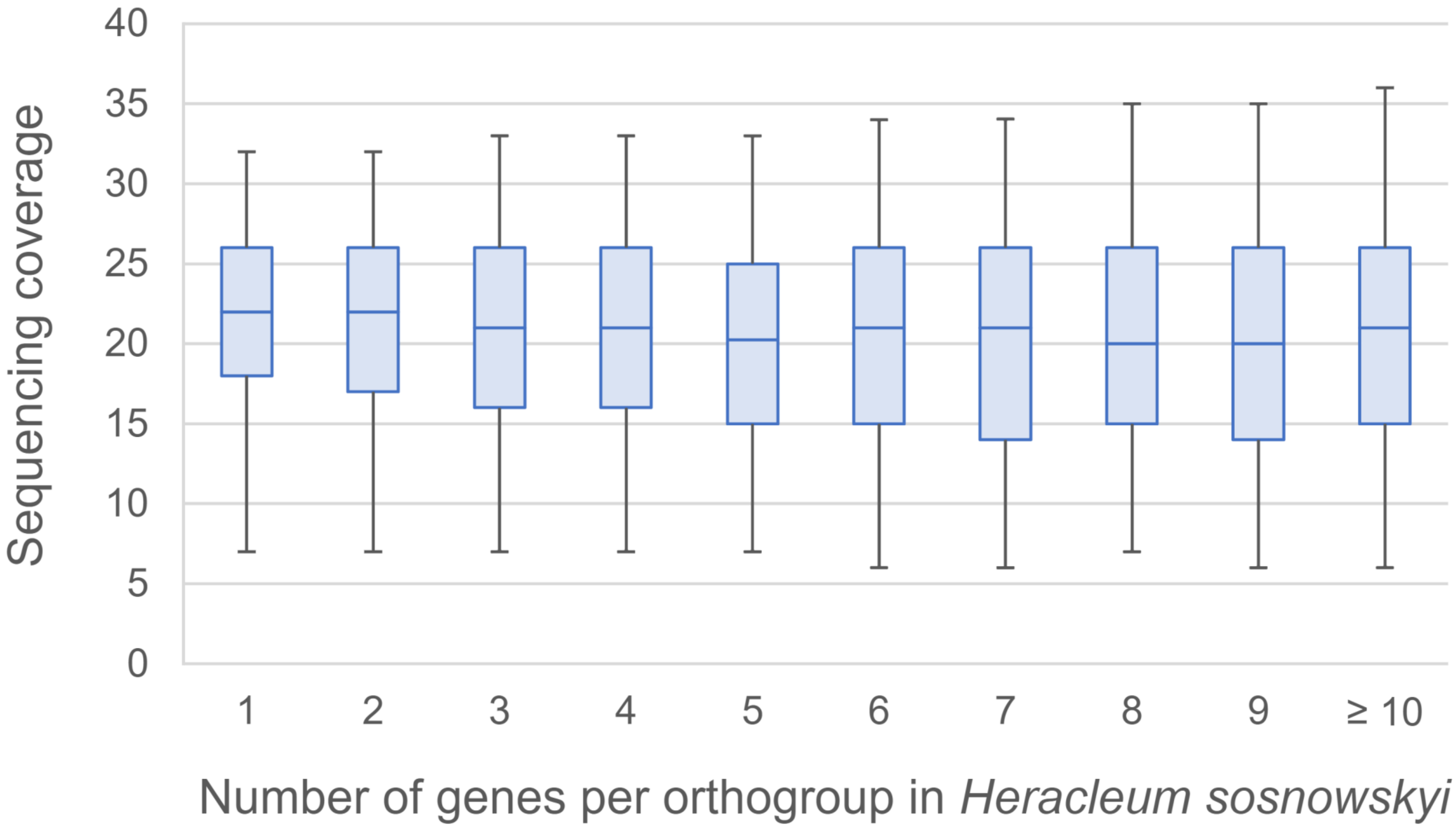


**Supplementary Figure 10.** Dependence between gene copy number per orthogroup and sequencing coverage in genes of *Heracleum sosnowskyi*. "Sequencing coverage" of a gene is the median coverage in the gene's exons by genomic reads. Box borders denote the 25th and the 75th percentiles, whiskers denote the 5th and the 95th percentiles, the middle line denotes the median.

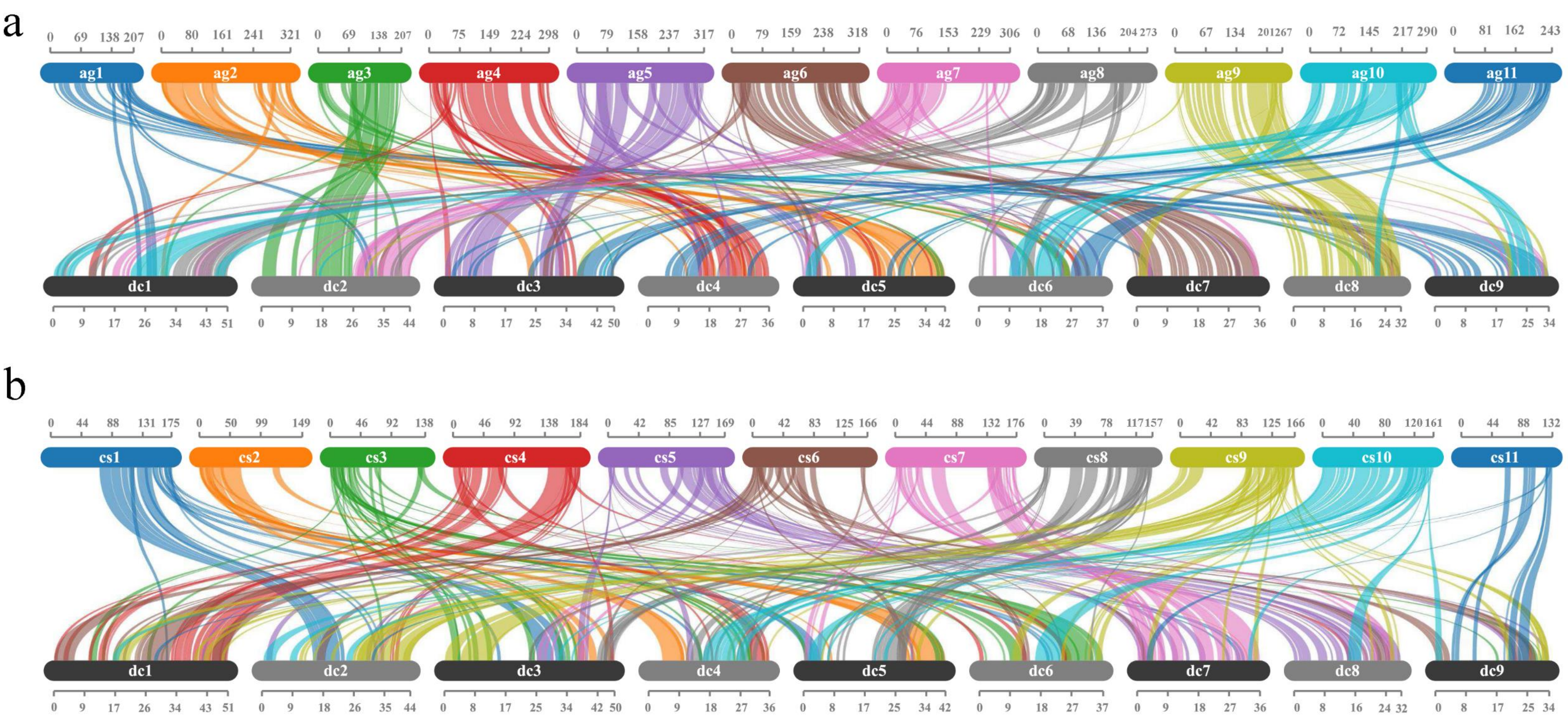


**Supplementary Figure 11.** Collinear blocks between genomes of Daucus carota and members of apioid superclade: *Apium graveolens* (a) and *Coriandrum sativum* (b). Only blocks that contain at least 20 pairs of homologous genes are shown. Coordinates are indicated in Mb.

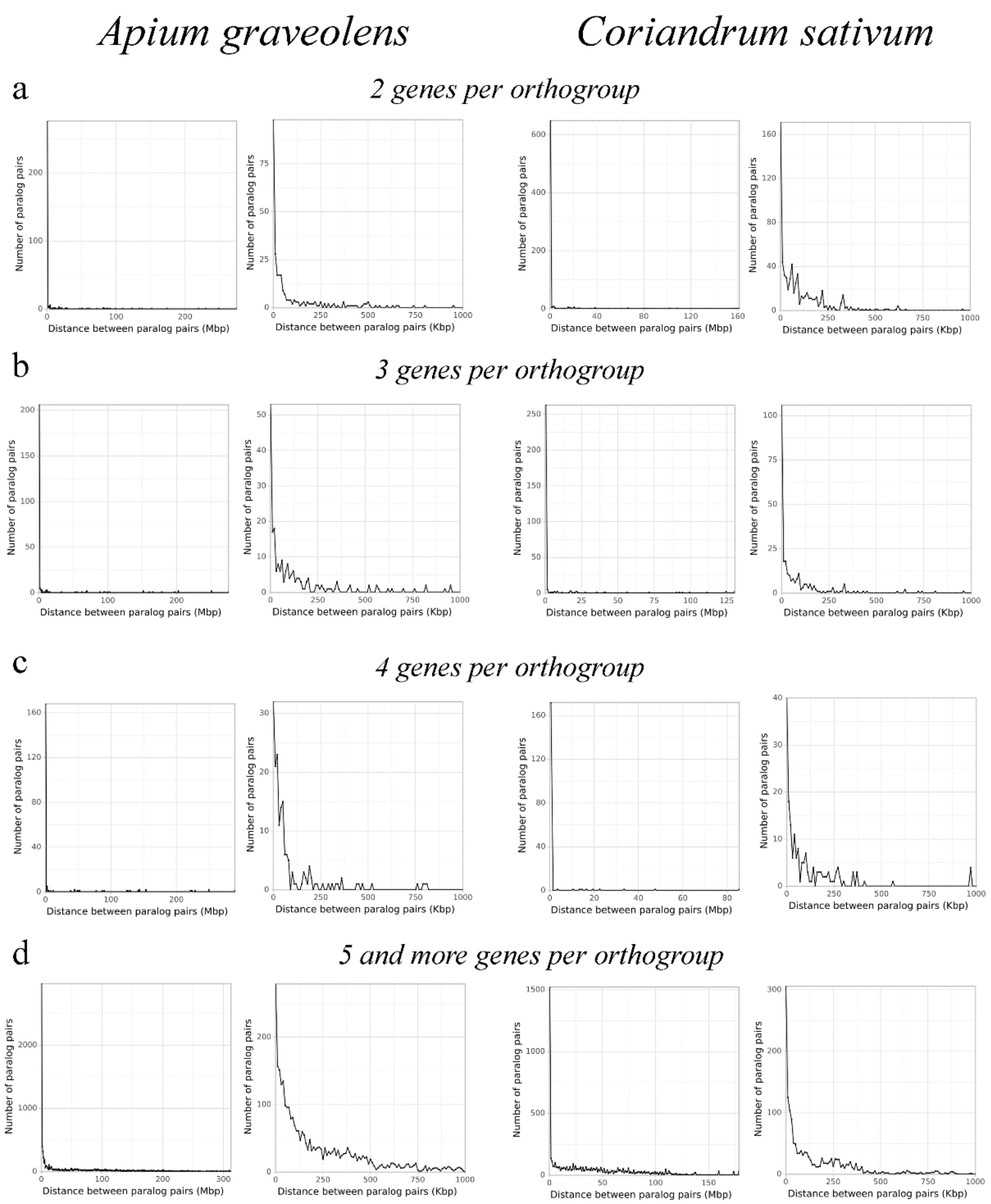


**Supplementary Figure 12.** Genomic proximity of duplicated genes in *Apium graveolens* and *Coriandrum sativum*.

a: distance between pairs of paralogs in orthogroups that have one *Daucus carota* and two *Apium graveolens*/*Coriandrum sativum* genes, left – whole range of variation of distances, 1-Mb bins, right – distances up to 1 Mb, 10-Kb bins.

b: distance between pairs of paralogs in orthogroups that have one *Daucus carota* and three *Apium graveolens*/*Coriandrum sativum* genes, left – whole range of variation of distances, 1-Mb bins, right – distances up to 1 Mb, 10-Kb bins.

c: distance between pairs of paralogs in orthogroups that have one *Daucus carota* and four *Apium graveolens*/*Coriandrum sativum* genes, left – whole range of variation of distances, 1-Mb bins, right – distances up to 1 Mb, 10-Kb bins.

d: distance between pairs of paralogs in orthogroups that have one *Daucus carota* and five or more *Apium graveolens*/*Coriandrum sativum* genes, left – whole range of variation of distances, 1-Mb bins, right – distances up to 1 Mb, 10-Kb bins.

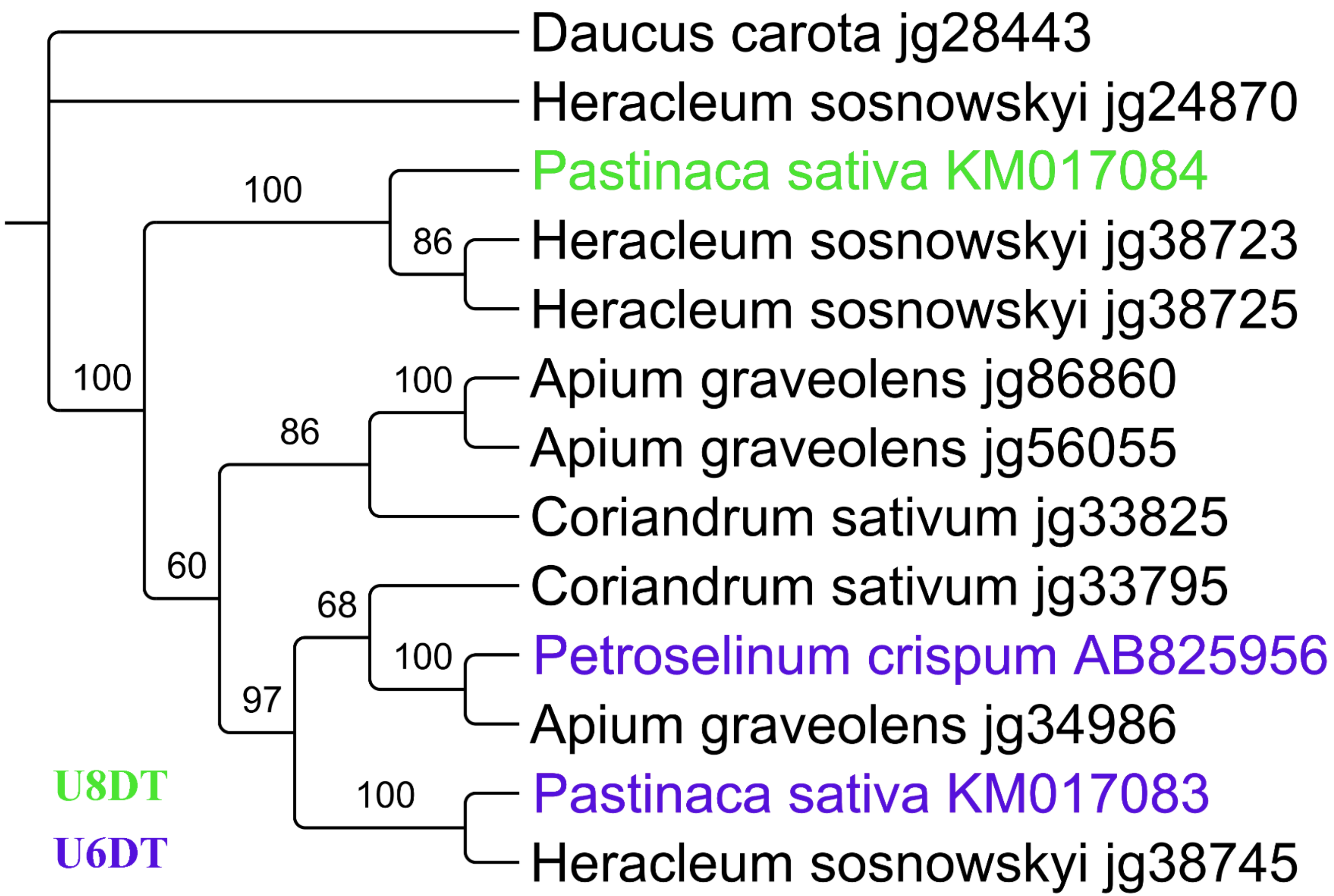


**Supplementary Figure 13.** Phylogenetic tree of UDT homologs in Apiaceae.

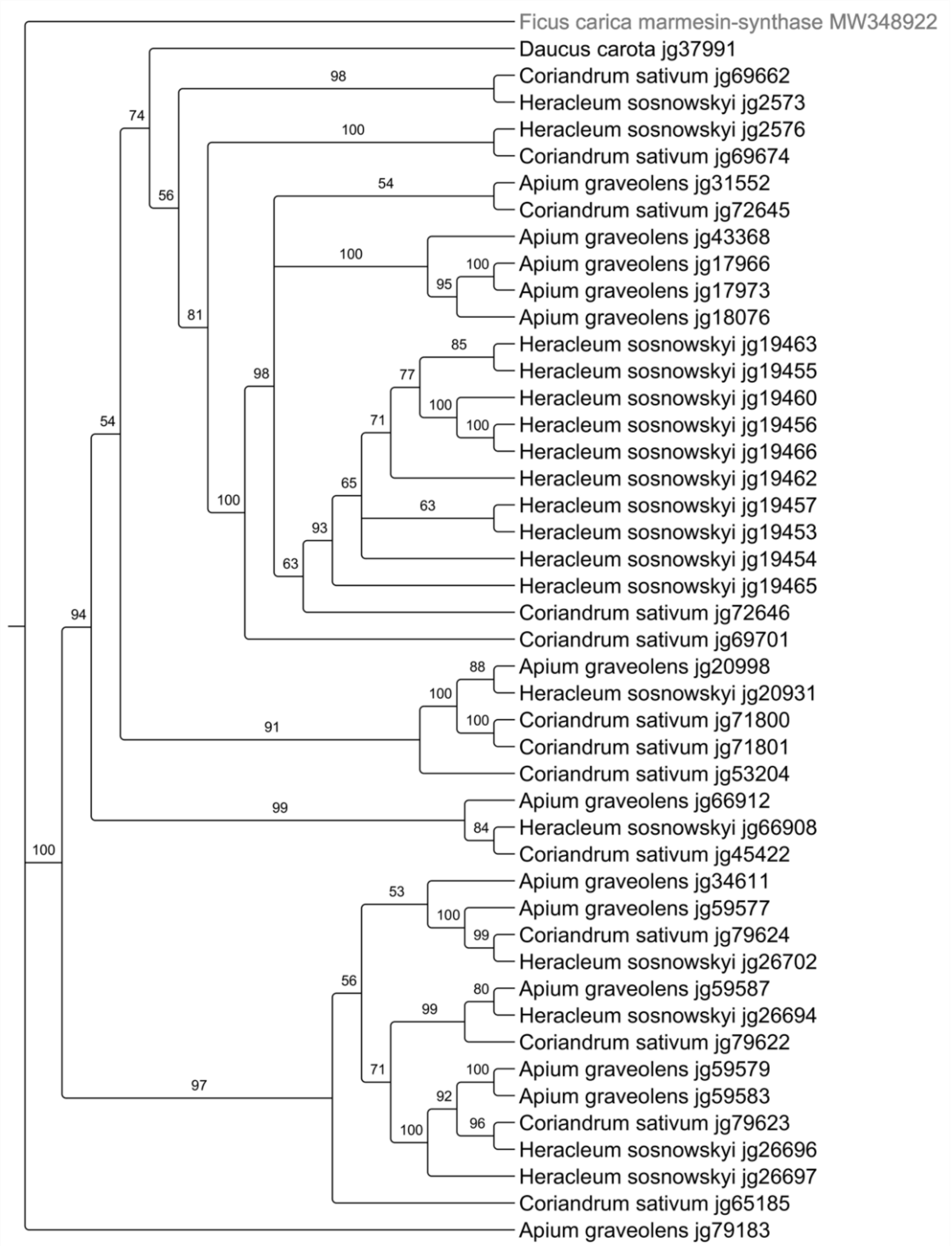


**Supplementary Figure 14.** Phylogenetic tree of marmesin-synthase homologs in Apiaceae

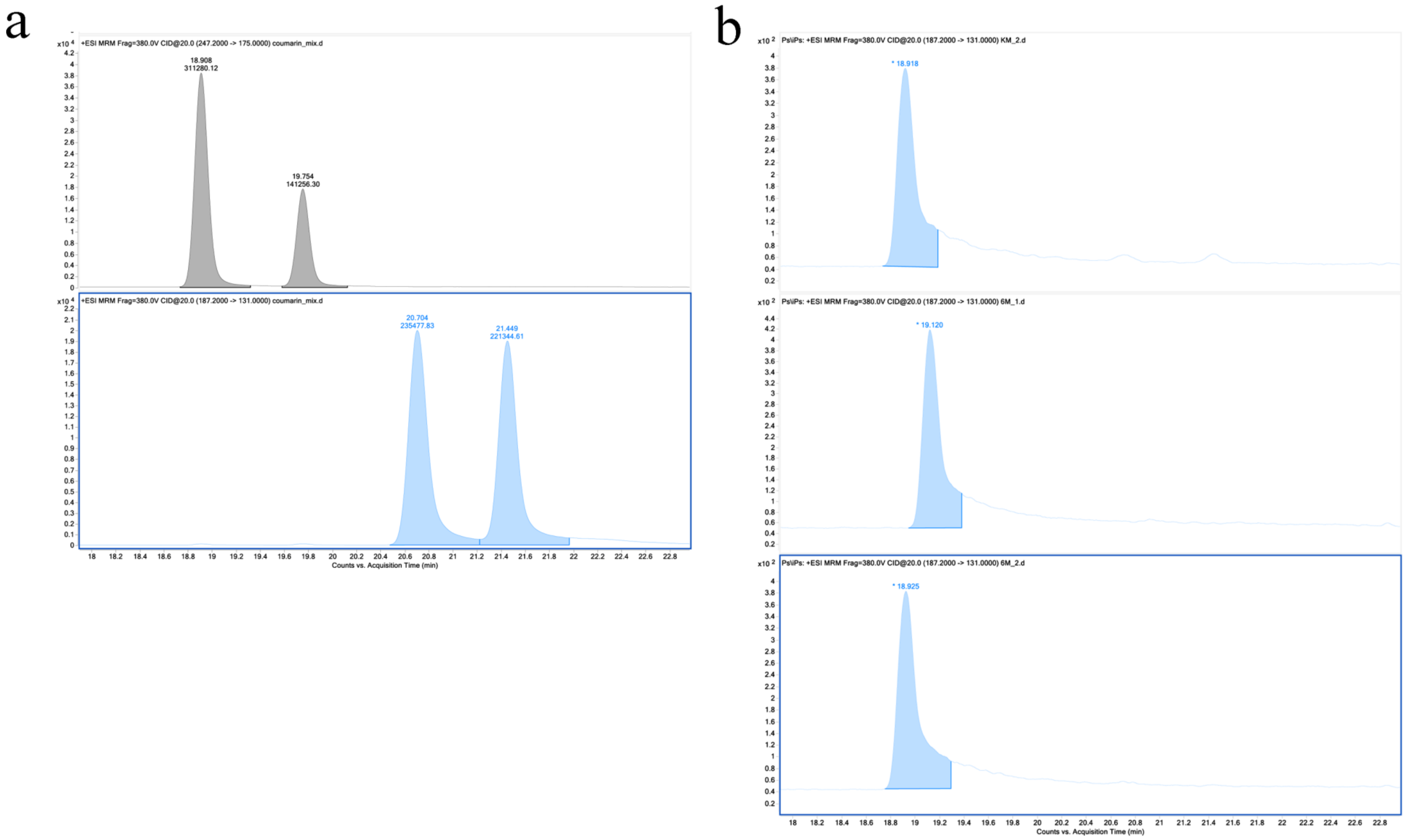


**Supplementary Figure 15.** Mass-spectrometry results for transgenic yeasts expressing jg70421 and jg70422 and control strain. (a) LC-MS/MS in MRM mode of the standard mixture consisting of marmesin ($t_R$ = 18.9 min), columbianetin ($t_R$ = 19.8 min), psoralen ($t_R$ = 20.7 min) and angelicin ($t_R$ = 21.4 min), (b) LC-MS/MS in MRM mode of the KM-2 (top), 6M-1 (middle) and 6M-2 (bottom) at the MRM transition of psoralen/angelicin. Slight (0.2 minute) shift in the retention times of the 6M-1 chromatogram is possibly due to the not completely thermally equilibrated column.

c

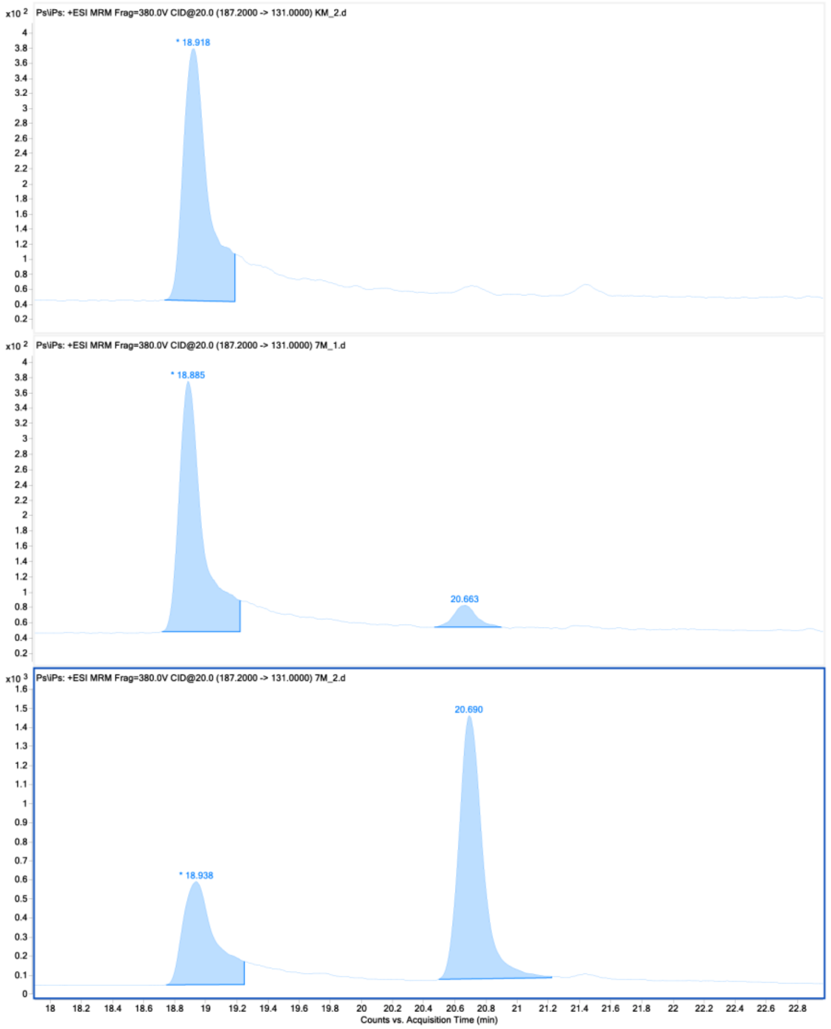


d

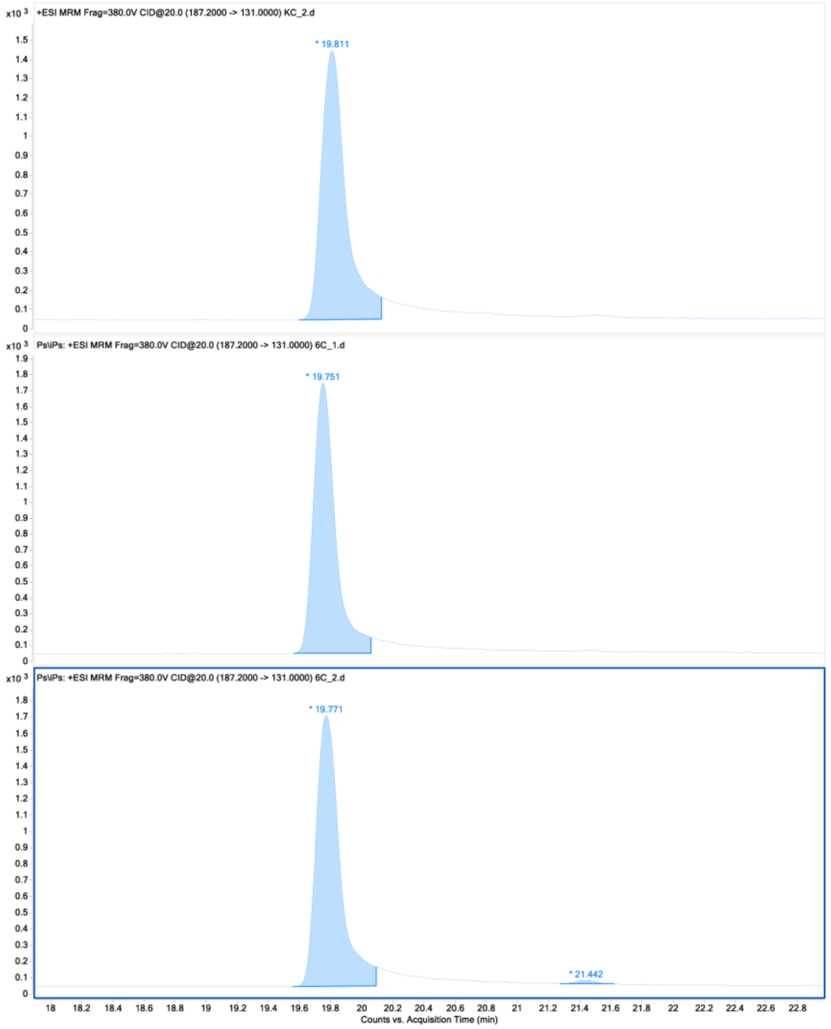


**Supplementary Figure 15 (continued).** Mass-spectrometry results for transgenic yeasts expressing jg70421 and jg70422 and control strain. (c) LC-MS/MS in MRM mode of the KM-2 (top), 7M-1 (middle) and 7M-2 (bottom) at the MRM transition of psoralen/angelicin, (d) LC-MS/MS in MRM mode of the KC-2 (top), 6C-1 (middle) and 6C-2 (bottom) at the MRM transition of psoralen/angelicin.

e

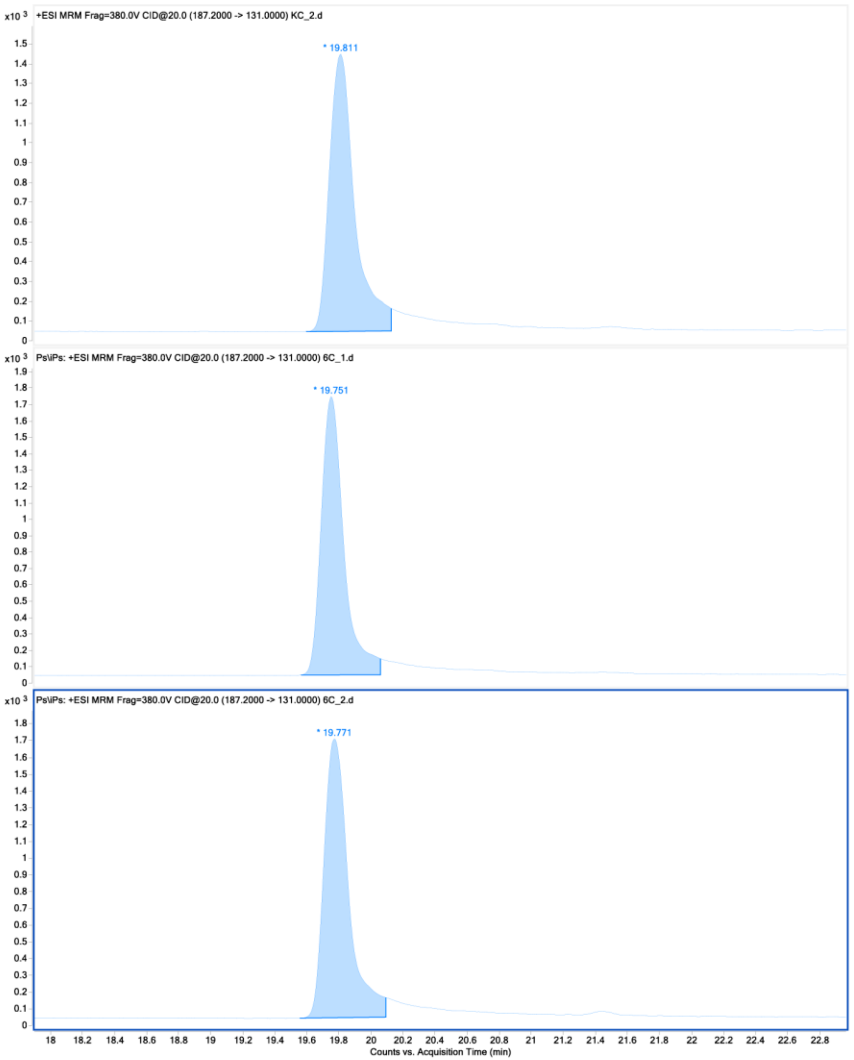


f

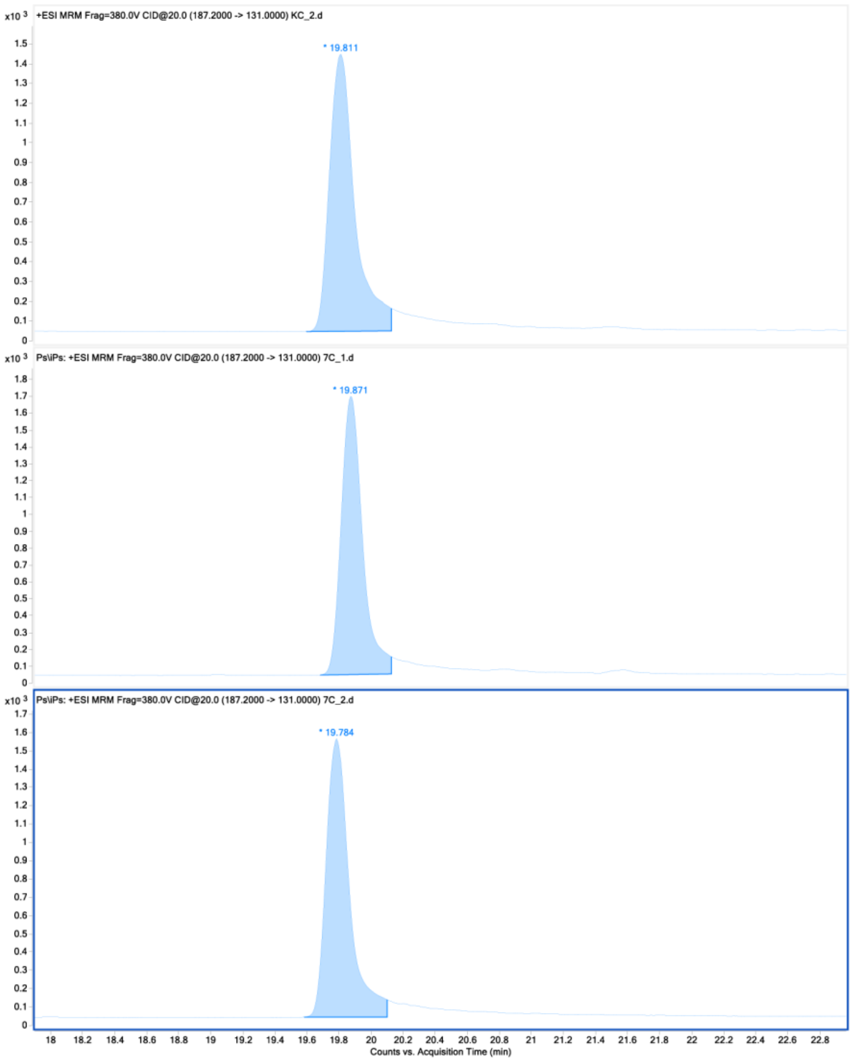


**Supplementary Figure 15 (continued).** Mass-spectrometry results for transgenic yeasts expressing jg70421 and jg70422 and control strain. (e) LC-MS/MS in MRM mode of the KC-2 (top), 6C-1 (middle) and 6C-2 (bottom) at the MRM transition of psoralen/angelicin (potential angelicin not shown), (f) LC-MS/MS in MRM mode of the KC-2 (top), 7C-1 (middle) and 7C-2 (bottom) at the MRM transition of psoralen/angelicin.
