## Supplemental experimental procedures for "The genome of the toxic invasive species *Heracleum sosnowskyi* carries an increased number of genes despite absence of recent whole-genome duplications"

**Nucleic acid extraction, library preparation and sequencing**

For genome sequencing (both WGS and Hi-C) we selected a single plant of *H. sosnowskyi* from the invasive area; the plant was collected in the wild and then grown in a greenhouse. DNA was extracted from leaves, freshly frozen in liquid nitrogen using CTAB protocol (Doyle and Doyle, 1987) and sequenced on Pacific Bioscience Sequel II platform (DNALink, South Korea). Nuclei for Hi-C (Lieberman-Aiden *et al.*, 2009) were isolated by gradient centrifugation using the CelLytic PN Isolation/Extraction Kit (Sigma-Aldrich, USA) according to the manufacturer's recommendations but using our modifications. First, leaf blades (without midrib) were homogenized with a blender in Nuclei Isolation Buffer (NIB). Then the homogenate (about 50 ml) was passed through several layers of a miracloth filter and centrifuged for 2 minutes at 100 g. Next, the supernatant was transferred into new 50 ml tubes and centrifuged for 10 min at 1200 g. We discarded the supernatant, and resuspended resulting pellet in 3 ml of NIB with the addition of 30 µl of Protease Inhibitor Cocktail for Plants (Sigma-Aldrich, USA) and Triton X-100 solution to a final concentration of 0.5%. Then the solution was applied to a gradient of sucrose (2.3 M) and 50% Percoll and centrifuged for 30 minutes at 3000 g. The fraction of nuclei located at the interface between the sucrose buffer and Percoll was pipetted and then purified by several rounds of centrifugation in NIB with the addition of a protease inhibitor. The resulting nuclei pellet was resuspended in 1 ml of NIB with the addition of formaldehyde (final concentration 1.5%). After 2 minutes of incubation, 75 μl of 2M glutamine solution was added to neutralize the formaldehyde. This step (cross-linking) is necessary to prepare Hi-C libraries from the nuclear fraction. The solution was centrifuged for 5 minutes at 2500 g. After removing the supernatant, the nuclear pellet was resuspended in 100 µl Nuclei PURE Storage Buffer and stored at –70°C. We used the EpiTect Hi-C Kit (Qiagen, Netherlands) to prepare Hi-C libraries. The procedures were held according to the manufacturer's protocol but using 30% fewer reagents. Library was checked for the length distribution and the absence of adapter dimers using capillary electrophoresis on Bioanalyzer2100 (Agilent, USA) and sequenced on Nextseq500 (Illumina, USA) using Nextseq High output 300 cycles kit in a paired-end mode. For transcriptome sequencing, we collected samples in two locations: one (the same where a sample for genome sequencing was collected) is near Moscow, the second is 1000 km to the North, in Karelia republic (see Supplementary table 1 for exact locations). Four above-ground structures were collected - leaves, flowers, inflorescence rays, and immature fruits. The samples were frozen on dry ice and used for RNA extraction. Before extraction, samples were homogenized in liquid nitrogen using mortar and pestle; for extraction we used RNEasy Mini kit (Qiagen, Netherlands) with the addition of Plant RNA Isolation Aid (ThermoScientific, USA) to the lysis buffer RLT. The quality of RNA was checked using capillary electrophoresis on Bioanalyzer2100 (Agilent, USA). 200-600 ng of high quality RNA was used for library preparation. At first step we selected polyadenylated mRNA using NEBNext Poly(A) mRNA Magnetic Isolation Module (New England Biolabs, USA). It was then fragmented for 5 minutes at 94 C and then processed with NEBNext UltraII RNA library preparation kit (New England Biolabs, USA) according to manufacturer’ instructions. Libraries were checked for the length distribution and the absence of adapter dimers using capillary electrophoresis on Bioanalyzer2100 (Agilent, USA) and sequenced on Hiseq4000 (Illumina, USA) using 300 cycles kit. Basecalling and demultiplexing of the data was done using bcl2fastq2 program (Illumina, USA).

For sequencing of RNA using long reads (Oxford Nanopore Technologies platform, ONT) we first converted RNA into double-stranded cDNA using Mint cDNA synthesis kit (Evrogen, Russia) and primers with custom barcodes. Then we pooled cDNAs representing different samples and prepared a library using ligation protocol (LSK-109 kit, Oxford Nanopore technologies, UK). Library was sequenced MinION instrument and flowcell v 9.4.1. Basecalling was done using Guppy 4.2.2 or Guppy 6.0.1 (Oxford Nanopore Technologies, 2019).

**Preprocessing of the reads**

No filtering was performed for PacBio HiFi reads. For ONT data reads with average Phred score below 7 were discarded during basecalling. Adapters were trimmed from Nanopore reads by Porechop 0.2.4 (Wick, n.d.) with default parameters. Illumina reads were trimmed by Fastp 0.21.0 (Chen *et al.*, 2018) by performing the following operations:

1.	Bases with Phred score below 3 were trimmed from 3' ends of reads.

2.	If there was a 5 bp-long region with average Phred score lower than 15, this region and everything towards the 3' end of the read were removed.

3.	If after the above procedures the average Phred score of a read was below 20, the read was discarded.

4.	If after the above procedures the length of a read was below 30 bp, the read was discarded.

5.	Adapters were trimmed using the default method of Fastp.

6.	If after the above procedures a read was discarded, its pair was discarded too.

**Genome assembly**

To achieve as accurate assembly as possible, the genome of *Heracleum sosnowskyi* was assembled by five different genome assemblers and then the assemblies were compared. The assemblies were performed by the following assemblers:

1.	Mabs-hifiasm 2.11 (Schelkunov, 2022) with default parameters. Both HiFi and Hi-C reads were used.

2.	Hifiasm 0.16.1 (Cheng *et al.*, 2021) with default parameters. Both HiFi and Hi-C reads were used.

3.	Flye 2.9 (Kolmogorov *et al.*, 2019) using HiFi reads, with default parameters.

4.	Miniasm 0.3 (Li, 2016) using HiFi reads, with default parameters. A PAF file with read overlaps, required by Miniasm, was made by Minimap 2.24 (Li, 2018) with parameters "-p 0.001 -N 999 -k19 -w19 -D --dual=no --no-long-join -e0 -m100 -I 1000G". Overlaps shorter than 1000 bp were removed. These options were chosen in order to achieve a good combination speed and accuracy of Miniasm, while keeping the size of the PAF file low.

5.	Canu 2.2 (Nurk *et al.*, 2020) using HiFi reads, with the option "genomeSize=1500m".

Removal of haplotypic duplications in all five assemblies was performed by Purge_dups 1.2.6 (Guan *et al.*, 2020) with default parameters.

Ten assemblies (five before Purge_dups and five after) were compared using the following metrics:

1.	Numbers of single-copy orthogroups, true multicopy orthogroups and false multicopy orthogroups, calculated by the program calculate_AG from Mabs 2.11 (Schelkunov, 2022). "False multicopy orthogroups" are orthogroups that have many genes because of assembly errors, while "true multicopy orthogroups" are orthogroups that actually contain many genes. The assembly with the largest numbers of single-copy orthogroups and true multicopy orthogroups and, at the same time, the smallest number of false multicopy orthogroups is the best.

2.	Results of BUSCO 5.3.2 (Simão *et al.*, 2015) analyses, performed using the "eudicots_odb10" dataset. Assembly with the largest completeness ("C") and the lowest number of duplicates ("D") was considered the best.

3.	N50, calculated by an in-house script. Assembly with the largest N50 was considered the best.

4.	Sum of contigs' lengths, calculated by an in-house script. The assembly with the length closest to the experimental estimate of 1750 Mbp (2C-value equal to 3.58 pg) (Zonneveld, 2019) was considered the best.

Based on the above criteria, the assembly made by Mabs without Purge_dups was considered the best (Supplementary Table 2). Scaffolding was performed for this assembly using Hi-C reads by Pin_hic 3.0.0 (Guan *et al.*, 2021) with default parameters.

To remove possible contamination, the scaffolds (here and further we call all sequences in the assembly "scaffolds", irrespective of whether they were scaffolds or contigs) were aligned to the NCBI nt database current as of 2022-04-20 by Megablast from BLAST+ suite 2.10.1 (Camacho *et al.*, 2009) with the maximum allowed e-value of 10^-5^. If more than half of the top 5 matches belonged to a taxon other than Embryophyta according to the NCBI Taxonomy database (current as of 2022-04-20) (Schoch *et al.*, 2020), the scaffold was considered as contamination. In the assembly of the genome of *H. sosnowskyi*, six scaffolds were determined as belonging to contamination, with a total length of 172,201 bp. Five of them were from bacteria and one from a fungus. When RNA or DNA reads were aligned to the assembly, contamination was always retained to minimize possible alignment of reads of the contamination to the genome of *H. sosnowskyi*. However, all information in the article (i.e., genome length and the number of genes) corresponds to the assembly after the removal of sequences of contamination, plastid sequences and mitochondrial sequences. To classify scaffolds into belonging to the nuclear, the plastid and the mitochondrial genomes, sequences of the plastid and the mitochondrial genomes should be determined. The plastid genome of *H. sosnowskyi* was assembled by GetOrganelle 1.7.5.3 (Jin *et al.*, 2020) with the option "-F embplant_pt". Since GetOrganelle 1.7.5.3 requires short reads for the assembly, we artificially made short reads by splitting HiFi reads into non-overlapping 600 bp-long regions. Each of these regions was, in turn, split into two 300 bp-long regions. This made artificial paired-end short reads, with lengths of 300 bp and the "insert sizes" of 600 bp. The plastid genome assembly was made using these reads. To assemble the mitochondrial genome of *H. sosnowskyi* we used the following technique, applied independently to contigs made by Mabs, Flye and Canu:

1) Mitochondrial genomes of Apiaceae available in the NCBI Nucleotide database as of 2022-05-24 were aligned by Blastn to contigs with the maximum allowed e-value of 10^-3^.

2) Average coverage by HiFi reads was calculated for all contigs in an assembly, by Minimap 2.24 and GATK 3.8.1 (McKenna *et al.*, 2010). Minimap run with default parameters was used for read alignment, while GATK was used to calculate the average read coverage based on this alignment.

3) A contig was considered probably mitochondrial if it had Blastn matches and, at the same time, had coverage by HiFi reads more than 50. 50 was chosen as the threshold because it is approximately two times higher than the coverage of nuclear contigs, which is approximately 25.

4) The genome assembly graph produced by a genome assembler was visualized in Bandage 0.8.1 (Wick *et al.*, 2015).

5) We tried to manually simplify the assembly graph of mitochondrial contigs into a structure consisting of single or several circles, with an additional requirement that sequencing coverage should be approximately equal in all regions of each circle.

The coverage of the longest mitochondrial contigs was approximately 400. Even if we discarded the contigs with coverage close 50 (they, potentially, belong to multicopy sequences transferred from the mitochondrial to the nuclear genome) we were still unable to simplify the mitochondrial graph into circles with uniform coverage. Additionally, we tried to assemble the mitochondrial genome by extending mitochondrial contigs by Elloreas 1.18 (Logacheva *et al.*, 2020). This assembly also didn't result in circles with uniform coverage. Finally, since we were incapable of assembling the mitochondrial genome of *H. sosnowskyi*, we considered all contigs determined using Blastn and sequencing coverage as described above as mitochondrial. Scaffolds that contained mitochondrial contigs were also considered mitochondrial.

To find plastid sequences in the assembly of *H. sosnowskyi* we aligned the plastid genome to the assembly by Blastn with the maximum allowed e-value of 10^-3^. If at least 50% of a scaffold length was covered by matches to the plastid genome, this scaffold was considered belonging to the plastid genome.

To find plastid and mitochondrial sequences in the assembly of *C. sativum* we aligned plastid and mitochondrial sequences of *C. sativum* from GenBank (accession codes MW477237.1, MW477238.1, KR002656.1) to the assembly by Blastn with the maximum allowed e-value of 10^-3^. If at least 50% of a scaffold length was covered by matches to the reference sequences, this scaffold was considered belonging to a plastid or a mitochondrial genome. For *A. graveolens* we used the same technique, but with plastid and mitochondrial sequences of this species as the reference (accession codes MK562756.1, MK036045.1). For *D. carota* we also used the same technique, but with plastid and mitochondrial sequences of this species as the reference (accession codes JQ248574.1, DQ898156.1).

Coupled with the technique used to find sequences of contamination this allowed us to classify scaffolds into belonging to the nuclear genome, to the plastid genome, to the mitochondrial genome and to the contamination for all of the four studied species of Apiaceae.

**Assembly validation and correction**

To find errors of assembly and scaffolding, the following three techniques were used:

1.	Short tandem repeats were searched for by TRF 3.0.9 (Benson, 1999) with the following parameters: "2 7 7 80 10 50 10 -h -d". Repeats with monomers TTTAGGG and CCCTAAA were considered telomeres. Telomeres not adjacent to an edge of a scaffold indicate a misassembly.

2.	Long repeats were searched for by RepeatModeler 2.0.3 (Flynn *et al.*, 2020) and RepeatMasker 4.1.2 (Smit *et al.*, n.d.). BuildDatabase (a tool provided with RepeatModeler) was run with the option "-engine rmblast" and RepeatModeler was run with the option "-LTRStruct". RepeatMasker was run with options "-e rmblast -xsmall -nolow -gff". Among the repeat families found by RepeatModeler and RepeatMasker was a family named "rnd-6_family-1721" which formed 11 long (on the order of megabases) tandem repeats with a monomer length of approximately 150 bp. Taking into account that *H. sosnowskyi* has 11 chromosomes, these likely are centromeric repeats. Presence of several centromeric repeats in one scaffold was considered an error.

3.	Alignment of Hi-C reads to scaffolds was visualized in Juicebox 3.1.4 (Durand *et al.*, 2016). Visually determined abnormal regions where adjacent regions had few connections by Hi-C reads indicate errors.

All errors that we found were errors of scaffolding and not errors of contig assembly, i.e., they could be fixed by cutting scaffolds in gaps ("NNN..." regions) and reconnecting the sequences in another order.

Based on the above-mentioned three criteria, we fixed the scaffolds by splitting them in gap regions and reconnecting the sequences in another order. After the reconnection, we visualized the Hi-C contact map by Juicebox again to verify that no close regions poorly connected by Hi-C reads remained.

**Genome annotation**

Prior to the structural annotation, repeats were found and masked by RepeatModeler and RepeatMasker as described above. To aid the structural annotation, Illumina and Nanopore RNA-seq reads were aligned to the genome. The Illumina reads were aligned by STAR 2.7.3a (Dobin *et al.*, 2013) with default parameters, while the Nanopore reads were aligned by Minimap 2.24 with the option "-x splice". Besides the RNA-seq reads, the structural annotation was aided by proteins of the "Plants" dataset from OrthoDB 10 (Kriventseva *et al.*, 2019). The structural annotation was performed by BRAKER2 (Brůna *et al.*, 2021), the version downloaded from GitHub by "git clone" on 2021-03-20, using the RNA-seq reads alignment and reference proteins, in the "ETP" mode. For the majority of genes BRAKER2 predicted a single isoform. For those with several isoforms, we removed all isoforms except the one with the longest CDS. False positive predictions (i.e., genes predicted by BRAKER2 that are actually not genes) were filtered by a number of methods:

1) Filtering by homology, method 1.

Proteins of the genes predicted by BRAKER2 were aligned by DIAMOND 2.0.15 (Buchfink *et al.*, 2021) to the NCBI nr (Sayers *et al.*, 2021) database current as of 2022-04-20 in the "--sensitive" mode with the maximum allowed e-value of 10^-5^. Genes whose proteins had matches were considered true positive predictions, i.e., actual genes.

2) Filtering by homology, method 2.

Proteins of the genes predicted by BRAKER2 were aligned by hmmscan from HMMER 3.3.2 (Mistry *et al.*, 2013) to the Pfam-A 35.0 (Mistry *et al.*, 2021) database with the maximum allowed e-value of 10^-5^. Genes whose proteins had matches were considered true positive predictions.

3) Filtering by high expression.

TPM values were determined for all genes by TPMcalculator 0.0.4 (Vera Alvarez *et al.*, 2019). The median TPM for genes that had matches in NCBI nr or Pfam (see "1)" and "2)") was calculated. Genes with TPM equal to or higher than this median were considered as "having high expression". All genes with high expression were considered true positive even if they didn't have matches in NCBI nr or Pfam. A gene may have high expression but no matches in NCBI nr or Pfam if has recently been *de novo* born (Van Oss and Carvunis, 2019) or if it accumulates mutations so quickly that it's hard to detect sequence homology with proteins from databases. For some genes, in particular ones related to the furanocoumarin biosynthesis pathway we performed manual annotation using the sequences of homologs from other Apiaceae. This was done by the BLAST alignment of CDS of these genes and the mark-up of exons based on this alignment.

All genes not supported by at least one of the above three criteria were considered false positives. In addition, even if a gene passed at least one of the above three criteria, it was considered a false positive if it met at least one of the following criteria:

4) The longest isoform of proteins encoded by a gene was shorter than 100 amino acids.

5) Among top 5 matches found by DIAMOND in NCBI nr more than half belonged not to Embryophyta according to the NCBI Taxonomy database current as of 2022-04-20. Such genes may belong to contamination.

6) Genes of transposable elements (TEs). Repeat masking by RepeatModeler and RepeatMasker was not 100% effective in masking TEs. Hence, an additional method for removal of TEs from the annotation was used. Descriptions, which are strings that describe proteins like "Floricaula/leafy-like transcription factor", were assigned to genes using the online version of PANNZER2 (Törönen and Holm, 2022) on 2022-10-05. A gene was considered belonging to a TE if descriptions of at least 10% of genes in its orthogroup (for how orthogroups were calculated see below) contained at least one of these substrings: "transpos" (it occurs as a part of strings like "retrotransposon" or "transposase"), "integrase", "copia", "gypsy", "gag-pol", "gag/pol", "gag-int-pol", "polyprotein", "RNA-directed DNA polymerase", "reverse transcriptase", "RNAse H", "ribonuclease H". The search for these substrings in gene descriptions was case-insensitive, i.e., irrespective of whether letters were uppercase or lowercase.

Here and further, we consider genes that had no orthologs as belonging to "singleton orthogroups", i.e., orthogroups that are composed of a single gene.

In addition to the structural (i.e., prediction of genes) annotation described above, a functional (i.e., description of genes' functions) annotation was performed, which consisted in assignment of gene descriptions and GO terms by PANNZER2.

For comparison with the genome of *H. sosnowskyi*, genomes of three other species of Apiaceae are used throughout the article. Namely, the genomes of *Coriandrum sativum* (Song *et al.*, 2020), *Apium graveolens* (Song *et al.*, 2021) and *Daucus carota* (Iorizzo *et al.*, 2016). A genome assembly and a genome annotation for *C. sativum* were downloaded from <http://cgdb.bio2db.com/Download/> . A genome assembly for *A. graveolens* was downloaded from NCBI GenBank (accession code GCA_902728035), while the annotation for *A. graveolens* is not publicly available. A genome assembly and a genome annotation for *D. carota* were downloaded from ENSEMBL (Cunningham *et al.*, 2022), release 53.

To evaluate the annotation quality for *C. sativum* and *D. carota*, we performed BUSCO analyses for genomes and for proteins from annotations and found that BUSCO's completeness for proteins was notably lower than for genomes, which indicates low quality of annotation. To fix this, we reannotated the genomes of *C. sativum*, *A. graveolens* and *D. carota* with exactly the same methods as we used to annotate the genome of *H. sosnowskyi*. During the annotation of the genome of *C. sativum* we used all RNA-seq reads available for this species in SRA as of 2022-07-02. The SRA identifiers of the datasets containing these reads are SRR8863732, SRR5410814, SRR5410771, SRR1700630, SRR1700873, SRR1700819. During the annotation of the genome of *A. graveolens* we used all RNA-seq reads available for this species in SRA as of 2022-07-02. The SRA identifiers of the datasets containing these reads are SRR14338343, SRR14338347, SRR14338344, SRR14338345, SRR14338346, SRR14338348, SRR14338349, SRR14338351, SRR11195970, SRR11195971, SRR11195972, SRR9104647, SRR9104648, SRR9104649, SRR9104650, SRR9104652, SRR9104651, SRR5575185, SRR5575184, SRR5575183, DRR003696, SRR1103267, SRR1023730, SRR14338350. As for *D. carota*, SRA contains many RNA-seq read datasets for this species due to its agricultural significance and a long-standing history of research. To accelerate the annotation, we used only reads from the article that described the assembly of the genome of this species (Iorizzo *et al.*, 2016). Datasets containing the reads have the following identifiers in SRA: SRR2148979, SRR2148980, SRR2148981, SRR2148982, SRR2148983, SRR2148984, SRR2148985, SRR2148986, SRR2148987, SRR2148988, SRR2148989, SRR2148990, SRR2148991, SRR2148992, SRR2148993, SRR2148994, SRR2148996, SRR2148997, SRR2148998, SRR2148999.

**Comparative genome analysis**

Orthogroups were determined for proteins of *H. sosnowskyi*, *C. sativum*, *A. graveolens*, *D. carota* and *Arabidopsis thaliana*. The proteins of *A. thaliana* were taken from the TAIR10 database (Berardini *et al.*, 2015). Orthogroups were calculated by OrthoFinder 2.5.4 (Emms and Kelly, 2019) using BLAST 2.10.1 as a tool for homology detection, MAFFT 7.450 (Katoh *et al.*, 2019) as a tool for multiple alignment and IQ-TREE 1.6.1 (Nguyen *et al.*, 2015) as a tool for phylogenetic tree construction (the options "-S blast", "-M msa", "-A mafft", "-T iqtree"). Synonymous substitution ratios ("dS") of paralog pairs were calculated by WGD 1.12 (Zwaenepoel and Van de Peer, 2019). Collinear blocks were found by MCScanX (Wang *et al.*, 2012), downloaded from GitHub using "git clone" on 2022-10-10. Collinear blocks were visualized by SynVisio online on 2022-10-18 (Bandi and Gutwin, Carl, 2020). Other analyses were performed by in-house scripts written in Python. Scripts are available on Figshare: **<https://figshare.com/articles/dataset/Genome_analysis_of_Heracleum_sosnowskyi_and_other_Apiaceae/21995738>**

**Analysis of furanocoumarin biosynthesis pathway genes and search for candidate genes for psoralen and angelicin synthase**

For the search of furanocoumarin biosynthesis pathway genes we used as query the sequences of the genes for which the function was shown experimentally, namely umbelliferone dimethylallyltransferases from *Pastinaca sativa* (accession numbers KM017083 and KM017084), marmesin synthase from *Ficus carica* (MW348922), psoralen-synthases from *Ammi majus* (AY532370) and *Pastinaca sativa* (EF191020) and angelicin-synthase from *Pastinaca sativa* (EF191021). For the genes of plants from Apiaceae we used 80% identity threshold and query coverage >90%, for one that are outside – 70% and 50% correspondingly. Sequences were aligned using MUSCLE (Edgar, 2004); phylogenetic analysis was performed using IQ-tree (Minh *et al.*, 2020). The analysis of amino acid residues important for protein function (substrate recognition sites, SRS) is based on the results of (Dueholm *et al.*, 2015) and (Larbat *et al.*, 2007).

**Expression of candidate genes in yeasts**

The cDNAs of Hsosn_jg70421 and Hsosn_jg70422 were amplified with gene-specific primers that included a restriction site for *EcoR*I and *Kpn*I: F-KpnI: GCAAAGGTACCAAAATGAAGATGCTGGAACAAAATCCC, R-EcoRI: AGCACGAATTCTCAAACATGYGGTGTGGCAAC using cDNA from leaf sample as a template. Separate amplification was not possible due to their high similarity, so amplification was performed using the same primers, and clones were separated after transformation based on the sequence. For heterologous expression of target genes, we used plasmids constructed on the base of the episomal shuttle vector pYeDP1/8-2 (further denoted as pYEDP) with Gal10-Cyc1 promoter, provided by Dr. D. Pompon (Center of Molecular Genetics, CNRS, Paris, France). This plasmid is capable to propagate in both *E. coli* and yeasts; it carries the ampicillin resistance gene and contains a yeast galactose-inducible promoter thus the production of a protein of interest is induced by galactose. The restriction enzymes EcoRI HF (New England Biolabs, USA) and KpnI (SibEnzyme, Russia) were used for the restriction of amplified cDNAs and vector simultaneously in NEB CutSmart buffer, with the addition of 1 µl of BSA for 1 hour at 37°C. Restriction results were analyzed using agarose gel electrophoresis; 7000 bp pYEDP vector fragment, and amplicon fragments (1500 nt) were excised from the gel and purified using Cleanup S-Cap kit (Evrogen, Russia) or DNA MinElute Gel Extraction Kit (Qiagen, Netherlands). Ligation was performed with Rapid DNA Ligation Kit (Thermo Scientific, USA) with a molar ratio of insert to vector in each reaction of 3:1 for 5 minutes at room temperature. The ligation mixtures were then purified using AMPure XP beads (Beckman Coulter, USA). Then *E. coli* cells XL1-blue strain (Evrogen, Russia) were transformed by electroporation. Electroporation was carried out using a Micropulser instrument (Bio-Rad, USA, protocol Ec1) in the SOC medium (SOB-medium with 0,2% glucose). After that, *E. coli* were incubated in a thermal shaker (1 hour, 37^o^C, 225 rpm), then they were plated on LB selection solid medium (LB-medium, Lennox) with ampicillin (100 ug/ml of medium) (AppliChem, Germany) . After 1 day, bacteria were transferred to a new plate after growing for one more day at 37 C, colonies were screened by PCR using Screen Mix (Evrogen, Russia) according to manufacturer instructions. Six positive clones were sequenced using Sanger sequencing on 3730 DNA Analyzer. Based on the obtained sequences, two clones of *E. coli* cells containing jg70421 or jg70422 were selected for further experiments. To isolate the plasmid with the desired insert, bacteria were grown ~~on~~ in liquid LB medium for a day (37 ^o^C, 225 rpm). Plasmids containing the target gene jg70421 or jg70422 were isolated using the Plasmid Miniprep kit (Evrogen, Russia) and were further used to transform yeast cells. In the present research, we used *Saccharomyces cerevisiae* strain 2805 (*MAT*α *pep4::HIS3, prb1_*δ*, can1,GAL2, his 3*δ*, ura 3-52*), kindly provided by Dr. S.-K. Rhee (GERI, Gyeongsangbuk-do, South Korea). Yeasts were inoculated into 5 ml of liquid YPD medium (1% yeast extract, 2% bacto-peptone, 2% glucose, pH 5.5) and grown with shaking (180 rpm) at 30°C overnight. Then, 1 ml of yeast suspension was inoculated into 5 ml of fresh liquid YPD medium, and cells were cultivated with shaking at 30°C for 4–5 hours up to OD600 = 1–2 (~ 2x10^7^ cells/ml). 6 ml of cell suspension was centrifuged (10 min, 4000 rpm, Eppendorf 5424, Germany), the sediment was washed with sterile TE buffer (with the resuspension of the cells), the cells were again pelleted by centrifugation (10 min, 4000 rpm), and the pellet was suspended in 200 µl of a sterile 0.1 M LiCl/TE solution. To transform the cells, 2–4 µl of plasmid DNA (0.5–2 µg) was added to 60 µl of the cell suspension, the sample was incubated for 6-8 minutes at room temperature, then to the sample was added 170 µl of a sterile solution 50% PEG-4000 in 0.1 M LiCl/TE. Further, the cells were incubated with shaking (50 min, 30°C, 150 rpm), 25 μl of DMSO was added and the cells were subjected to heat shock (5-8 min at 42°C). After adding 1 ml of sterile TE buffer (10 mM Tris-HCl, pH 8.0, 1 mM EDTA) to the sample, the cells were thoroughly suspended and precipitated by centrifugation (5 min, 4000 rpm). The pellet was suspended in 200-300 µl of sterile TE buffer and rubbed into a plate with selective SD medium (0.67% yeast nitrogenous bases, 2% α-D glucose, 0.1% casamino acids or 0.5% (NH4)_2_SO_4_., 1.5-2% agar). To grow the transformed clones, the plates were incubated at 30°C for 5 days. The expression of recombinant proteins in yeast cells was performed according to Akiyoshi-Shibata and coworkers (Akiyoshi-Shibata *et al.*, 1991). Recombinant strains were cultivated in a liquid YPD medium with shaking (180 rpm) at 30°C overnight, then 1 ml of culture was diluted with 20 ml of a liquid synthetic SD minimal medium and the cell suspension was grown under the same conditions for 24 hours. Synthesis of the heterologous proteins in transformed yeast cells was induced by replacing glucose with galactose in the culture medium: the cells were washed twice with 10 ml of sterile water and resuspended to a final OD600 ~ 1.0 in 20 ml of a selective SG induction medium containing 2% galactose (0.67% yeast nitrogenous bases (Yeast Nitrogen Base, Difco), 2% α-D galactose, 0.1% casamino acids, or 0.5% (NH4)_2_SO_4_). Expression of heterologous genes in *S. cerevisiae* cells was performed for 18-44 hours (30°C, 180 rpm). Induction of heterologous gene expression was checked by qRT-PCR with primers specific to the insert: 70421-22-F TTGATGCTTTTCTGGAAGGT and 70421-22-R: GAATCTCAAGCAAAATGGATA. To test the functionality of the cloned genes, 4 hours after the transfer of recombinant yeast cells to 20 ml of SG culture medium, the suspension was divided into 2 parts of 10 ml. Psoralen precursor marmesin (final concentration 100 uM) was added to one part, and angelicin precursor columbianetin (final concentration 100 uM) was added to the second part. After cultivation for 20 and 44 hours, 5 ml of suspension were taken. Suspension samples were centrifuged (4000 g , 5 min), then the supernatant was mixed with methanol in a ratio of 1:1 and used for mass-spectrometry analysis in order to detect the presence of the expected compounds (psoralen and angelicin for marmesin and columbianetin as precursors, respectively. For each gene the analysis was run in two biological replicates.

**Targeted LC-MS/MS analysis**

The LS-MS/MS measurements were performed using the equipment of “Human Proteome” Core Facilities of the Institute of Biomedical Chemistry (Russian Federation). 0.5 ml of the sample was centrifuged for 30 min at 20,000 rpm, then 100 µl of the supernatant was transferred to a clean vial and evaporated to dryness in a vacuum concentrator (Vacuum concentrator plus, Eppendorf, Germany) at 45°C. Samples were re-dissolved in 20 μl of methanol and subjected to analysis. Agilent 1290 Infinity LC system was used in connection with an Agilent 6495 Triple Quad mass spectrometer (Agilent technologies, USA) with an electrospray ionization source.

Mobile phases were prepared with the following materials: water purified with a Milli-Q Integral 3 system (Millipore, France), formic acid (ThermoFisher Scientific, USA), acetonitrile (ThermoFisher Scientific, USA). Composition of the mobile phases were: mobile phase A - MilliQ water with 0.1% formic acid, mobile phase B - acetonitrile 0.1% formic acid. Injuction volume of 1 µl was used in all cases.

Chromatography was performed using Waters Acquity UPLC BEH C8, 1.7 µm, 2.1 x 100 mm column with a Waters Acquity UPLC BEH C8 1.7 µm 2.1 x 12.5 guard column (Waters Corporation, USA) at the flow rate 0.25 ml/min, and column temperature of 40°C. Gradient elution conditions were as follows: isocratic elution at 5% of mobile phase B and 95% of mobile phase A until the 3 minute mark, followed by the linear gradient to 30% of mobile phase B at 18 minutes, then another linear gradient to 100% of mobile phase B at 19 minutes with isocratic wash until the 22 minute mark. Then within a minute the solvent composition was switched back to 5% of B followed by a column equilibration step until the 50 minute mark. MS detection parameters: positive ionization mode, source temperature 300°C, carrier gas pressure 4.8 bar, drying gas flow rate 12 l/min, capillary voltage 4 kV. Samples were analyzed using MRM mode, the ion transitions were as follows: for psoralen and angelicin - ion-precursor - 187.2 m/z, ion product - 131.0 m/z, for marmesin and columbianetin - ion-precursor - 247.2 m/z, ion product 175.0 m/z. Collision energy was 20 eV for all substances.
